# *FICTURE:* Scalable segmentation-free analysis of submicron resolution spatial transcriptomics

**DOI:** 10.1101/2023.11.04.565621

**Authors:** Yichen Si, ChangHee Lee, Yongha Hwang, Jeong H. Yun, Weiqiu Cheng, Chun-Seok Cho, Miguel Quiros, Asma Nusrat, Weizhou Zhang, Goo Jun, Sebastian Zöllner, Jun Hee Lee, Hyun Min Kang

## Abstract

Spatial transcriptomics (ST) technologies have advanced to enable transcriptome-wide gene expression analysis at submicron resolution over large areas. Analysis of high-resolution ST data relies heavily on image-based cell segmentation or gridding, which often fails in complex tissues due to diversity and irregularity of cell size and shape. Existing segmentation-free analysis methods scale only to small regions and a small number of genes, limiting their utility in high-throughput studies. Here we present FICTURE, a segmentation-free spatial factorization method that can handle transcriptome-wide data labeled with billions of submicron resolution spatial coordinates. FICTURE is orders of magnitude more efficient than existing methods and it is compatible with both sequencing- and imaging-based ST data. FICTURE reveals the microscopic ST architecture for challenging tissues, such as vascular, fibrotic, muscular, and lipid-laden areas in real data where previous methods failed. FICTURE’s cross-platform generality, scalability, and precision make it a powerful tool for exploring high-resolution ST.

## Introduction

Spatial transcriptomics (ST) has important implications for elucidating cell-cell interactions and localizing specific activities in biological/pathological processes^1,2^. ST can be divided into sequencing-based and *in situ* imaging-based technologies^3^. First, sequencing-based ST annotates each molecule with spatially resolved barcodes and performs RNA-seq to profile the entire transcriptome. However, initial sequencing-based ST^4^, now commercialized as 10X Visium, is largely limited by its low resolution (>100 µm). Recent developments have dramatically improved the resolution of spatial barcodes through microfluidics-based (DBiT-Seq^5^; down to 20 µm), bead-based (Slide-Seq^6^, HDST^7^; down to 4 µm)^5^, and next-generation sequencer (NGS)-based (Seq-Scope^8^, Stereo-Seq^9^, and Pixel-Seq^10^; down to 0.5 µm) approaches. In particular, the latest NGS-based ST approaches achieve submicron resolution comparable to an optical microscope. Second, *in situ* imaging-based ST reveals the location of specific transcripts through multiplexed molecular profiling using *in situ* hybridization or *in situ* sequencing. These high-resolution (<0.5 µm) technologies based on optical microscopy typically assay hundreds of genes using combinatorial encoding as implemented in commercial platforms such as Vizgen MERSCOPE^11^, 10X Xenium^12^, and CosMx SMI^13^.

In line with these rapid developments, enormous amounts of submicron-resolution ST datasets are being generated on an unprecedented scale. One challenge is that, even if billions of transcripts are captured per tissue section, typically only a few transcripts are observed per µm^2^, and conventional algorithms have limited accuracy when analyzing at micron-resolution. For these reasons, published analyses of submicron-resolution ST datasets aggregate individual transcripts into either fixed-size grids or image-based cell segments. Image-based cell segmentation has been considered the gold-standard when nuclear staining such as DAPI or cell boundary staining is available. After cell segmentation, various methods such as Squidpy^14^, Giotto^15^, Seurat^16^, or GraphST^17^, can be applied for downstream inference.

However, image-based cell segmentation faces a number of practical challenges in analyzing real-world datasets: (1) Because three-dimensional tissue is processed into two-dimensional sections, nuclei could be missed for many cells while their cytoplasmic RNAs is still captured (Fig. 1A). This leads to incorrect assignment of transcripts to adjacent cells. (2) Irregularly shaped cells are often inaccurately segmented by existing methods that assume a spherical shape around the nucleus. (3) Cells without nuclei (e.g., red blood cells) or multinucleated cells (hepatocytes, muscle, etc.) skew cell segmentation. (4) Cell boundary markers may not be expressed at all in certain cell types. (5) Both cell size and cell density vary dramatically even within a single tissue type. (6) Variations in sample quality and processing can hinder the acquisition of high-quality reference images sufficient for single cell segmentation. (7) Cell segmentation ignores extracellular RNA, which may be vital for some biological processes such as cell-cell communication^18,19^. Indeed, we observed that 12-55% of transcripts are dropped during image-based cell segmentation in published ST datasets (Fig. 1B). Therefore, even with state-of-the-art cell segmentation algorithms or manual cell segmentation by trained pathologists, accurate analysis of submicron-resolution ST datasets is challenging and often infeasible with image-based cell segmentation.

**Figure 1.**
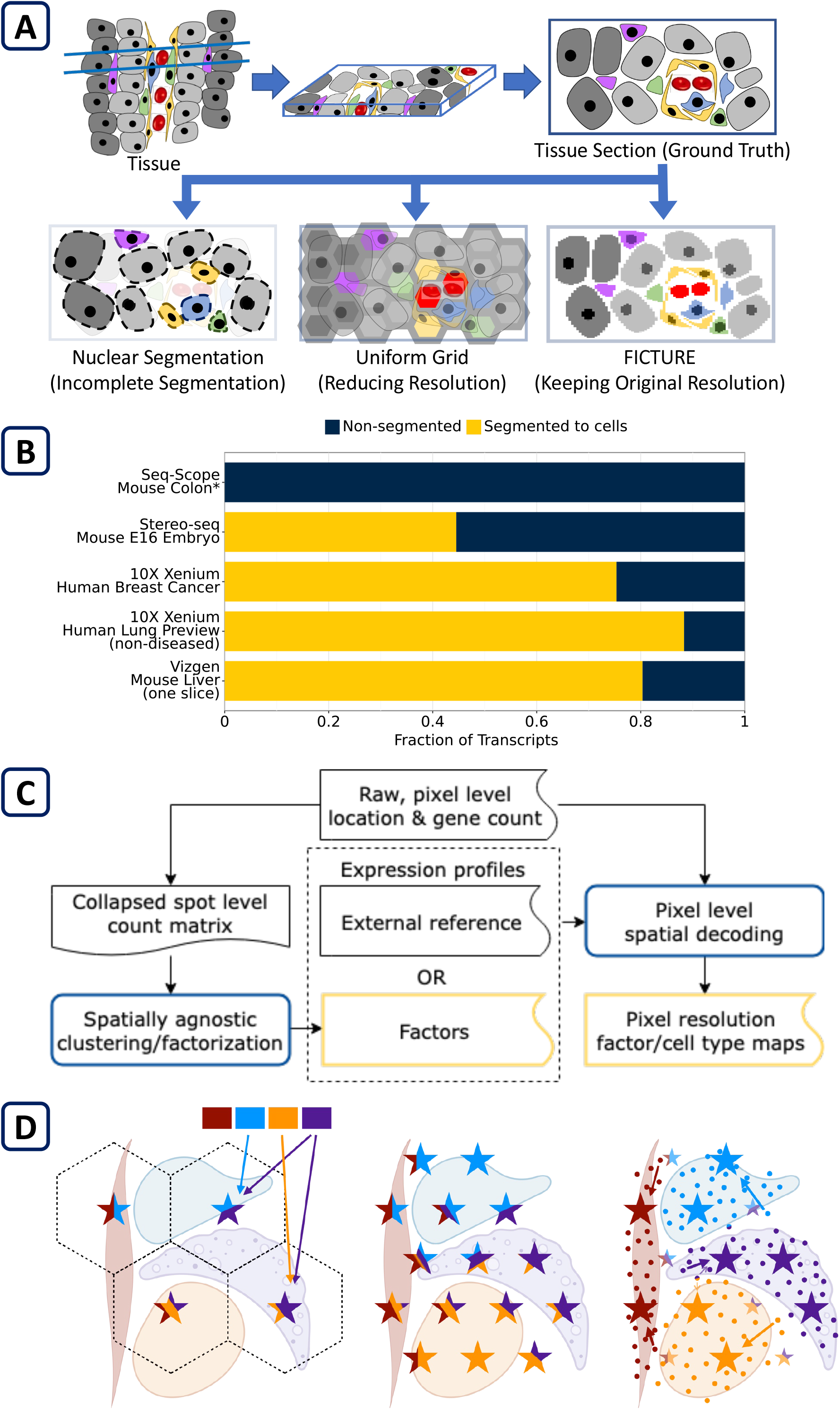
Overview of *FICTURE*. (A) Illustration of the ST analysis on a two-dimensional (2D) section of three-dimensional (3D) tissue. The 2D view of each cell varies substantially depending on the shape of the cells and the location and orientation of the cutting plane. A 2D slice may not capture some nuclei, leading to bias in nuclear segmentation based on histological staining (bottom left, dashed lines). Uniform gridding substantially compromises the original resolution (bottom center). FICTURE preserves the original resolution at the pixel level (bottom right). (B) The proportion of transcripts included (yellow) or excluded (navy) in the cell segmentation analysis across five high-resolution datasets used in this study. Asterisks (*) indicate that Seq-Scope dataset was not segmented into cells. (C) Schematic illustration of the FICTURE’s workflow. FICTURE’s pixel-level inference is based on factor-specific expression profiles. These can be generated from (i) unsupervised factorization using Latent Dirichlet Allocation (LDA) on spot level gene counts collapsed according to a hexagonal grid (default; D, left), (ii) other tools that provide clustering or factorization (e.g. Seurat, scanpy, squidpy) on collapsed data, or (iii) external sc/snRNA-seq reference with cell type specific gene expressions. (D) Schematic illustration of the FICTURE’s algorithm. Based on the initial factors, FICTURE places anchors on a lattice denser than cells (D, center) and infers a mixture distribution over factors at each anchor based on the gene expressed at pixels in its neighborhood. Each pixel is assigned to a nearby anchor probabilistically and the pixel’s sparse gene expression is modeled by that anchor’s mixture distribution. Initially pixels are assigned to the nearest anchors deterministically, then the anchors’ mixture distributions over factors and the pixel-to-anchor assignments are updated iteratively. Upon convergence, pixels are assigned to anchors with factor mixtures best explaining the pixels’ gene expression; and anchors tend to collect information from a more homogeneous set of pixels (D, right).

Because of these limitations, previous studies have explored alternative approaches. One of the simplest methods is to partition the ST data into fixed-size grids. Most published sequencing-based ST datasets are not paired with nuclei or cell boundary staining, which is required for imaged-based cell segmentation. Consequently, these datasets were analyzed with uniform grids^8,9^, which typically captures multiple cells in a single grid, obscuring the signature of small or spatially dispersed cell types. Grid segmentation also compromises the submicron resolution of the raw data, and often fails to accurately delineate cell type boundaries (Fig. 1A, lower center). Adaptive segmentation algorithms, such as the volume-distance-based segmentation algorithm, aim to dynamically segment the ST dataset based on transcript densities^10^. While this was effective in analyzing Pixel-Seq brain data, it is difficult to generalize to many other tissue types where various cell types are densely packed together. Sliding window algorithms, as implemented in SSAM^20^ and STtools^21^, partially mitigates the loss of resolution during segmentation, while the oversampling of transcripts limits the choice of clustering algorithms.

Segmentation-free techniques for analyzing high-resolution ST data are a promising alternative (Fig. 1A, lower right), but existing methods have limited scalability. One of the most accurate methods, Baysor^22^, uses Markov Random Field to assign individual transcripts to clusters without relying on external segmentation, but it does not scale to larger tissue areas (>5mm^2^) or to transcriptome-level gene panels (>1,000 genes), as later examined in this study. To the best of our knowledge, there are no scalable segmentation-free inference methods that can analyze the vast amounts of data produced by cutting-edge submicron-resolution ST technologies.

Here we present FICTURE (Factor Inference of Cartographic Transcriptome at Ultra-high REsolution), a segmentation-free spatial factor analysis method that makes inference on individual spatial coordinate points (i.e., pixels) at submicron resolution. Our method scales to the whole transcriptome (i.e., >20,000 genes) and billions of pixels associated with transcriptional readouts and. FICTURE can learn the factors unsupervised or take user-provided reference expression profiles to construct a pixel-resolution spatial map of such profiles. We have applied our method to data generated by four state-of-the-art high-resolution ST platforms, including NGS-based Seq-Scope and Stereo-seq, and *in situ*-based MERFISH and Xenium. Our method reveals cellular heterogeneity at a microscopic scale, precisely delineates cell type boundaries, and identifies rare cell types from their surroundings. It also eliminates blind spots of cell segmentation in challenging tissues such as skeletal and smooth muscle, tumor stroma, adipose tissue, and tissues with complex vasculature, outperforming existing methods. To our knowledge, FICTURE is the only segmentation-free algorithm that can perform microscopic resolution analysis on both NGS-based and *in situ*-based ST data while efficiently scaling to the largest ST datasets currently available.

## Results

### Segmentation-free factor inference at submicron resolution

FICTURE aims to reconstruct the fine-scale tissue structure by first decomposing gene expression patterns across the tissue section into factors then using local information to assign each pixel to these factors probabilistically. Each factor defines the average expression levels of all genes, and we model the molecule counts generated from a factor using a multinomial distribution (Supplementary Fig. 1A). From a biological perspective, these factors may represent cell types, cell functions, or cell states specific to certain physiological or pathophysiological conditions; they may even represent subcellular or extracellular transcriptomic phenotypes. FICTURE by default learns factors unsupervised from the dataset by aggregating pixels into hexagonal grids to fit a standard Latent Dirichlet Allocation (LDA) model assuming independence between the hexagons (Supplementary Fig. 1B), similar to existing methods for low-resolution data^23^. Alternatively, the factors can also be obtained from external sc- and sn-RNA-seq references datasets, or from other spatially agnostic methods such as *Seurat*^16^ and *Scanpy*^24^ that learn factors based on grids. FICTURE can use these external factors as priors to initialize the LDA factors or use them directly as input factors (Fig. 1C).

To reconstruct pixel-level tissue architecture with these factors, FICTURE adaptively aggregates the extremely sparse information from pixels locally using anchor points to infer the latent factors at each pixel without deterministic segmentation. Specifically, FICTURE defines anchors as lattice points over the tissue region and lets each anchor learn a probability distribution over the factors (Fig. 1D). FICTURE updates pixel and anchor level parameters iteratively. Each pixel picks an anchor from its neighborhood probabilistically then samples the gene counts according to the chosen anchor’s mixture distribution. Each anchor in turn aggregates information from the pixels in its neighborhood to update its mixing probabilities across factors. Since the information is shared only locally, FICTURE constructs inference units as conditionally independent spatial patches (500 µm-sided squares by default) for efficient parallel processing within each minibatch. Upon convergence, FICTURE delineates cell types unconstrained by the lattice (Fig. 1D, right) so that inferred pixel-level factors represent single cell transcriptome architectures within the tissue.

We applied FICTURE to simulated datasets, as well as five real ST datasets assayed with four different state-of-the-art submicron resolution platforms. These real datasets consist of a mouse colon assayed with NGS-based Seq-Scope, a mouse E16 whole embryo assayed with NGS-based Stereo-seq, human breast cancer and healthy lung datasets assayed with *in situ*-based 10X Xenium, and a mouse liver dataset assayed with *in situ*-based Vizgen MERSCOPE. (Supplementary Fig. 2).

### FICTURE reveals microscopic tissue architecture in Seq-Scope colon data

To illustrate the advantage of pixel-level factor inference over existing approaches, we first applied FICTURE to a mouse colon dataset generated by the updated Seq-Scope^HISEQ^ technique, which provides a larger field of view than the original Seq-Scope^MISEQ^ ^8^. We performed biopsy-induced injury of the mouse colonic mucosa and analyzed healing wounds 24 hrs after injury (red-outlined arrows in Fig. 2A, J. K)^25,26^. The data contains 6.8 million spatial barcodes from 9 tissue sections jointly covering ~18mm^2^ area (Supplementary Fig. 3A), and one representative section is presented in Fig. 2B. This dataset is accompanied by H&E histology from the same section; however, the dataset is not amenable to image-based cell segmentation due to the extreme irregularity and diversity in cell morphology associated with different cell layers.

**Figure 2.**
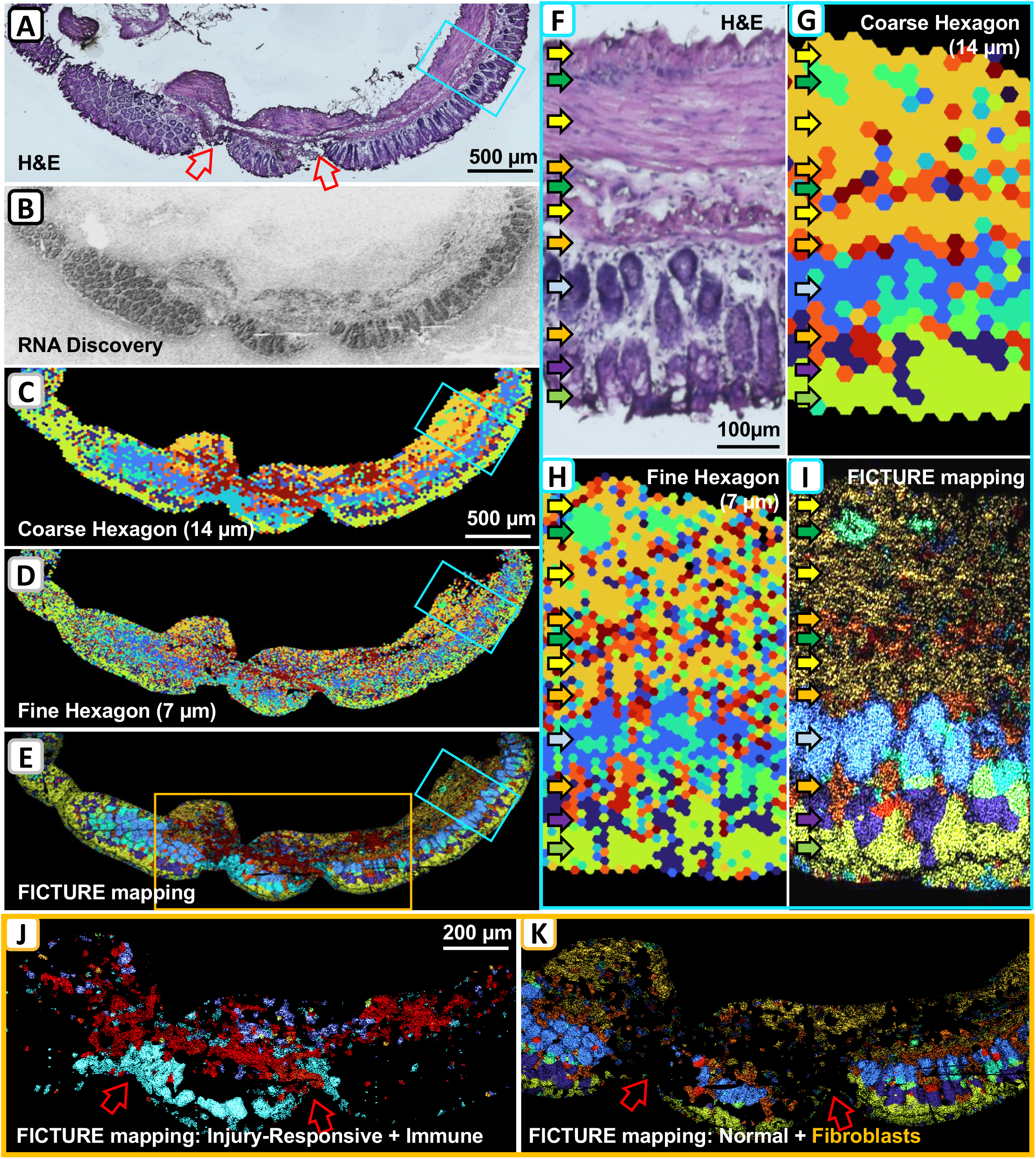
Grid-based vs. pixel-level analysis of Seq-Scope mouse colon. Mouse colon sections with biopsy-induced injuries were profiled with Seq-Scope, segmented into hexagons with side length 14 µm, and clustered with Seurat, similar to the published study^8^. These clusters are provided as input factors for spatial factorization in three different resolutions. Nine sections were analyzed (Supplementary Fig. 3), and one is shown here. Wound sites are indicated by red arrows in (A), (J) and (K). (A) Hematoxylin and eosin (H&E) staining. (B) Density of transcripts observed at each pixel. (C) Cluster assignments of the original 14 µm-sided hexagons. (D) Projection of the clusters onto 7 µm-sided hexagons. (E) Pixel-level decoding using FICTURE. (F-I) Close-up view of blue rectangles in (A), (C), (D) and (E), showing normal colonic wall layers in H&E (F), hexagonal clusters (G, H), and FICTURE inference (I). Arrows indicate muscle layers (yellow), ganglion cells in myenteric and submucosal plexus (green), fibroblasts in submucosa and lamina propria (orange), crypt epithelial cells (light blue), goblet cells (purple), and surface colonocytes (light green). (J) Visualization of clusters corresponding to injury-response and immune cell infiltration near the injury site. (K) Visualization of normal colon cells and fibroblasts in the same region. The full color codes and marker genes of each factor are shown in Supplementary Table 1.

The data were initially partitioned into 43,494 14 μm-sided hexagons and clustered using *Seurat*. These clusters revealed cell-type diversity in normal colon, as well as injury-responsive epithelial and immune cell populations (Supplementary Fig. 3B). However, these cell types’ spatial distributions are too coarse (Fig. 2C) to capture the spatial complexity shown in the histology (Fig. 2A). Making the grid smaller slightly improved the resolution but led to extremely noisy cluster assignments due to the low expression counts in each grid unit (Fig. 2D). In contrast, the pixel-level factor inference by FICTURE dramatically improved the resolution without introducing noisy artefacts (Fig. 2E) for all examined tissue sections (Supplementary Fig. 3C). Higher magnification image shown in Fig. 2F-2I demonstrates that the FICTURE can identify all layers of the uninjured colonic wall. FICTURE identified the longitudinal and circular smooth muscle layers of the muscularis propria and the muscularis mucosae (yellow), ganglion cells in myenteric and submucosal plexus (green), fibroblasts in the submucosa and lamina propria (orange), epithelial crypts (light blue), goblet cells (purple), and surface colonocytes (light green) (Fig. 2F, I; each layer marked by arrows). These details were far less evident in grid-based analyses (Fig. 2G, H).

FICTURE was also able to identify the cell phenotypes associated with mucosal injury (Fig. 2J, 2K) with molecular details that are not discernible by imaging-based histological examination. For instance, FICTURE was able to detect infiltration of different immune populations into the injury site. We also found distinct injury responses of epithelial and smooth muscle layers, involving different immune cell populations, which were difficult to detect by the histological examination. Although some of these features were also appreciable through grid-based analysis, only FICTURE was able to associate the transcriptome feature with underlying histopathological findings with high resolution precision.

### Simulation study assesses FICUTRE’s accuracy, robustness, and scalability

To evaluate the accuracy and the scalability of FICTURE’s pixel-level spatial factor inference, we compared it with two existing unsupervised methods, Baysor^22^ and GraphST^17^. Baysor was the only pixel-level inference method available for comparison, and GraphST was selected as a representative of methods based on cell segmentation. We simulated data based on ten selected cell types from a scRNA-seq reference dataset^27^, assigning 8 cell types to specific regions and scattering cells from the other 2 cell types throughout the space to mimic infiltrating cells (Fig. 3A). Due to Baysor’s limited scalability with increasing number of genes, we only selected 500 highly variable genes for analysis. Since GraphST requires segmented data, we provided the true simulated cell boundaries for GraphST but not for the other methods.

**Figure 3.**
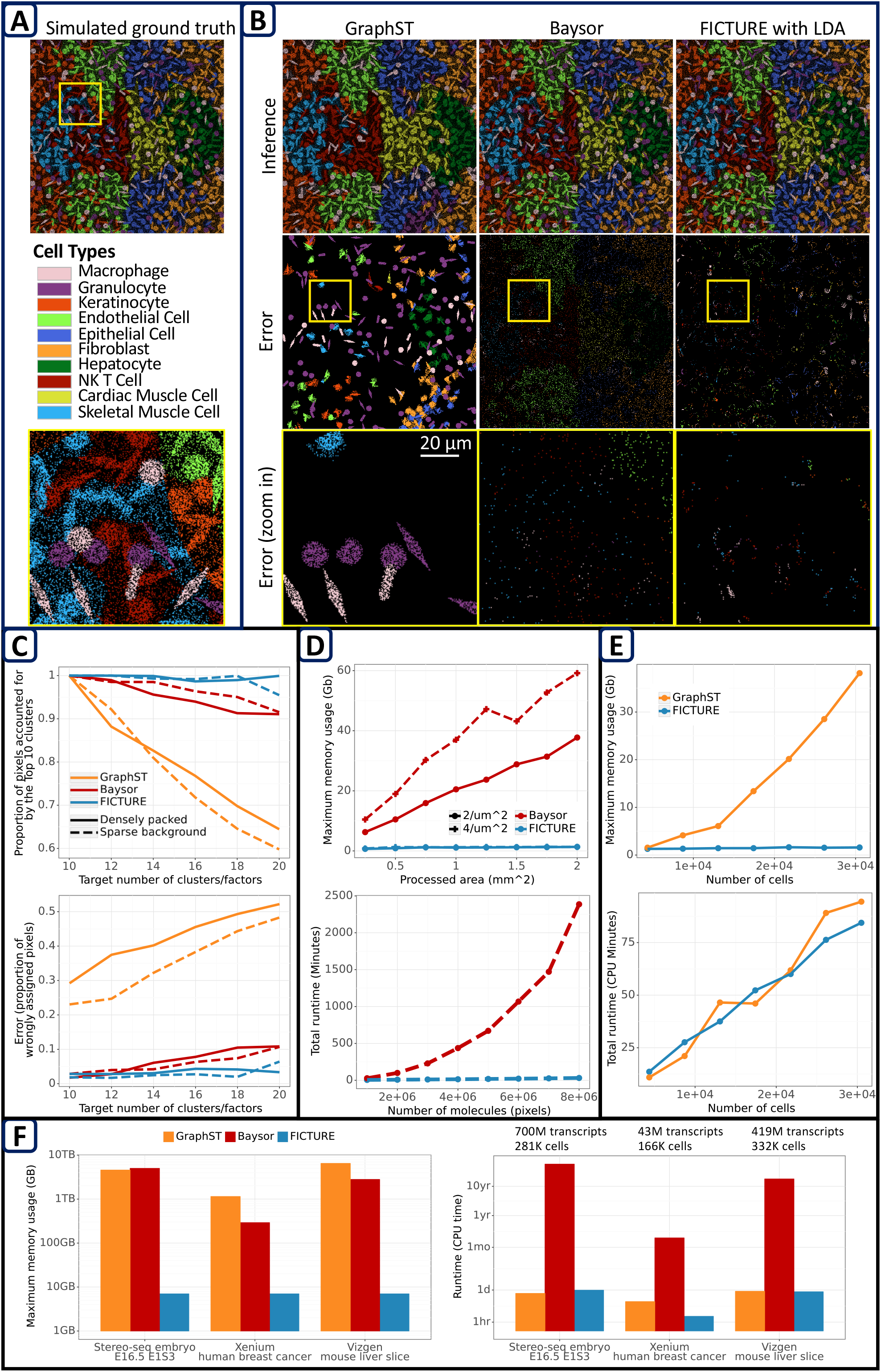
Comparison between FICTURE, Baysor, and GraphST using simulation. (A) The simulated ground truth with 10 cell types, which are colored as indicated. The top panel shows the full region, and the bottom panel shows the close-up view of yellow-boxed region (100 µm x 100 µm). (B) Comparisons between GraphST (left), Baysor (center), and the fully unsupervised FICTURE initialized by LDA (right). The top row represents pixel-level cluster assignment; the middle row represents the difference between inferred and true clusters; the bottom row magnifies the same area as in the bottom panel of (A). (C) Robustness to over-specified number of clusters when the true number of clusters is 10. Metrics are quantified as the coverage of the pixels by top 10 factors (top) and pixel-level assignment errors (bottom) of GraphST, Baysor, and FICTURE. Densely packed data are presented in Supplementary Fig. 4, while sparse background data are presented above in (A) and (B). (D) Comparisons of segmentation-free methods (Baysor vs. FICTURE) in terms of memory consumption (top) and running time (bottom) as a function of the area processed (mm^2^) or the number of molecules (pixels). The top and bottom figures are based on the same set of experiments. (E) Comparison with segmentation-based methods (GraphST) in terms of memory consumption and running time as a function of the number of cells (although FICTURE is performed at the pixel level). (F) Projected memory consumption and computational time to perform the analysis on real datasets using each method. Due to the limited scalability of Baysor, the projection assumes that only 500 genes are used in Stereo-seq.

Since all three methods require user-specified number of clusters, we first evaluated these algorithms when the correct number of cell types (k=10) is given. Both FICTURE (98%) and Baysor (97%) demonstrated high accuracy at pixel level, whereas GraphST tended to be less accurate (78%) especially for scattered cell types due to oversmoothing (Fig. 3B, 3C). Similar results were obtained when simulation data with more densely packed cells and transcripts were used (Fig. 3C, Supplementary Fig. 4). In both analyses, FICTURE and Baysor accurately delineated cell type boundaries for both region-specific and scattered cell types, regardless of cell shapes.

Next, we evaluated the robustness of these methods under an over-specified number of clusters. This aspect is critical because, in most cases, we do not have prior knowledge of how many factors or cell types are present in the given sample. In the above simulation with 10 cell types, across model sizes from 10 to 20 clusters/factors, only a small fraction of pixels were assigned to the extra factors in FICTURE (<4.5%) and Baysor (<8.9%), demonstrating their high robustness (Fig. 3C). On the other hand, GraphST tends to over-cluster to achieve the target number of clusters and assignied a large fraction (20~50%) of pixels to the extra factors. This is similar to observations from other graph-based methods that use community detection algorithms such as Louvain^28^ and Leiden^29^.

Computational scalability of analysis method is a crucial factor as high-resolution high-throughput datasets accumulate. FICTURE operates on a fixed memory budget even with increasing data size since FICTURE performs parallel processing in minibatches. In contrast, both Baysor and GraphST need to store the full data in memory. Baysor’s memory consumption scales quadratically with the number of genes and linearly with the number of molecules (Fig. 3D); GraphST’s memory consumption scales super-linearly with the number of cells (Fig. 3E). As a result, the memory footprint of FICTURE is orders of magnitude smaller than the other methods and the difference will increase as the cutting edge technologies generate datasets with larger field of view and/or larger number of genes (Fig. 3D, E). FICTURE’s runtime scales linearly with the number of pixels, while Baysor’s runtime scales quadratically with the number of molecules, regardless of the molecule density (Fig. 3D). For example, to analyze a 1 mm^2^ area at 4 transcripts/µm^2^ with 500 genes, Baysor requires 37GB of memory and 7.3 CPU hours while FICTURE requires 1.2GB of memory and 0.23 CPU hours. GraphST, which runs at the cell level, requires 1.5GB and 0.18 CPU hours with runtime scaling linearly with the number of cells per iteration. We also estimated the required resources and evaluated the accuracy of these three methods in real datasets (Fig. 3F), as we detail at the end of the section.

### Precise, scalable and unbiased profiling of the whole transcriptome of a mouse embryo

To evaluate FICTURE on a real-world high-resolution ST dataset with a large field of view, we analyzed the whole transcriptome profiling of a mouse E16.5-stage sagittal section by Stereo-Seq^9^, which contains 221 million pixels with a total of 700 million transcripts in a 115 mm^2^ region^9^. FICTURE used 8.7 μm-sided hexagons to fit the unsupervised LDA model with 48 factors and then assigned individual pixels to factors, with the top 21 (30) factors accounting for >95% (99%) of the pixels. We compared the spatial distribution of the 25 published cell type annotations derived from DAPI-based nuclear segmentation followed by graph-based clustering^9^ with the pixel-level factors inferred by FICTURE independently of the DAPI images (Fig. 4A-4C). DAPI-based nuclear segmentation removed a majority (55.5%, 389 million) of transcripts from the analysis. The proportion of dropped transcripts varies by tissues, from 34.3% in pancreas to 94.0% in blood vessel (Supplementary Fig. 5A-5F). For example, in the blood vessels, erythrocytes were systematically underrepresented in the DAPI-based segmentation compared to FICTURE’s pixel level inference because fully differentiated red blood cells lack nuclei (Supplementary Fig. 5A-5F). The proportion of dropped transcripts also varies by genes. Not only hemoglobin genes but also many other heart- or muscle-enriched genes (*Myl7, Tnni3, Myh6, Myl3, Tnni2, Cox8b, Myl4*) show high proportion (>60%) of dropped transcripts; overall, more than 17,500 (59%) genes had more than half of the transcripts dropped (Supplementary Fig. 6A), suggesting widespread loss of molecular information during cell segmentation.

**Figure 4.**
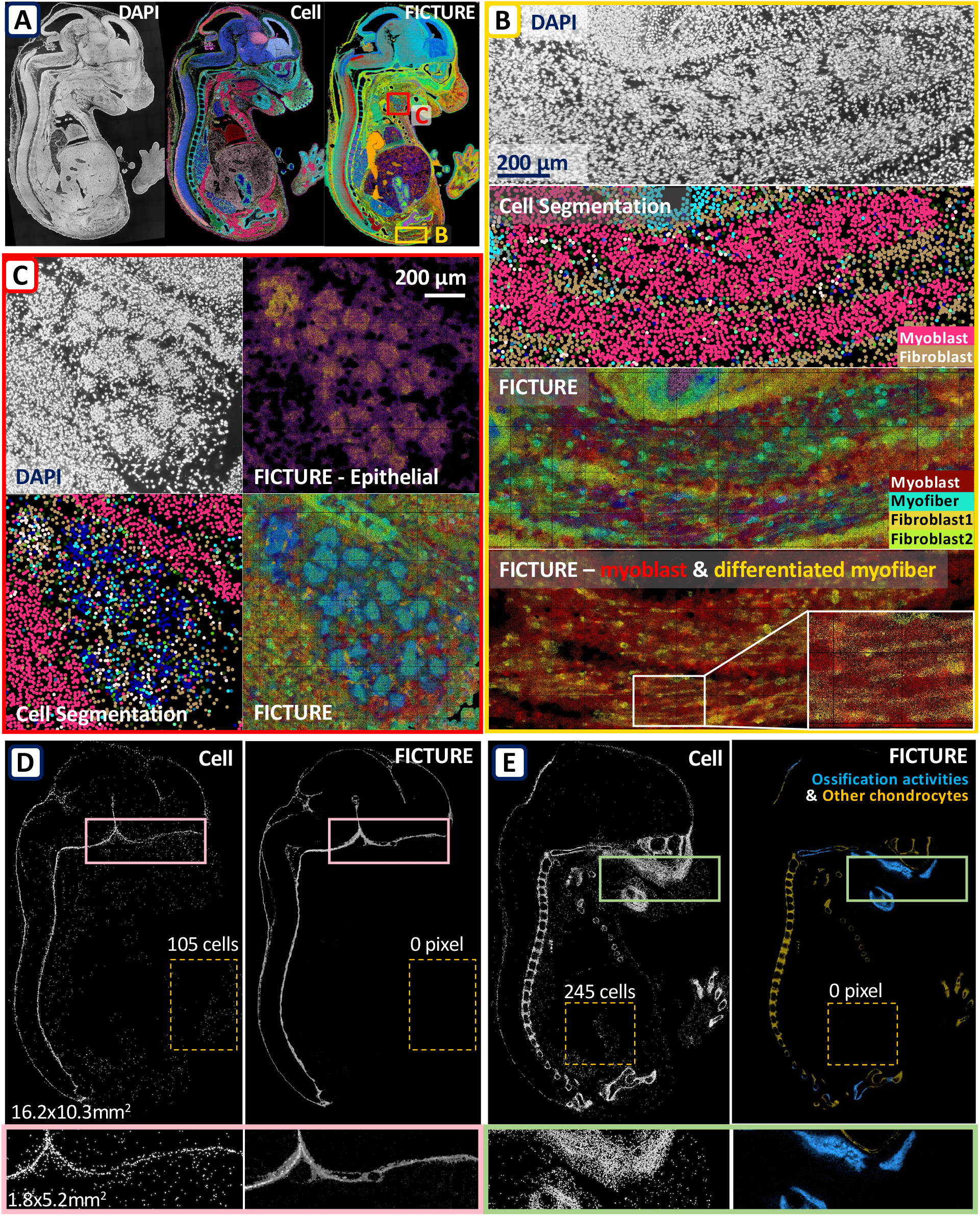
Application of FICTURE to a whole mouse embryo dataset by Stereo-seq. (A) The published E16 mouse embryo Stero-seq dataset was used to produce following images. Left: DAPI staining image. Center: published analysis based on cell segmentation. Right: pixel-level analysis using a fully unsupervised FICTURE. (B) Yellow-boxed area was magnified focusing on developing skeletal muscle. From top to bottom: DAPI staining image, cell-based segmentation image from the MOSTA browser, pixel-level decoding with FICTURE, and two FICTURE factors representing myoblasts (red) and differentiating myofibers (yellow). (C) Magnification of the submandibular gland. The upper left corner is the DAPI image. The lower left corner shows the cell-based segmentation visualized at the pixel level. The lower right corner shows the pixel-level decoding with FICTURE. The upper right corner shows the posterior probability of an epithelial factor at each pixel. (D) Spatial distribution of meningeal cell types from the published cell segmentation analysis (left), and pixel-level inference with FICTURE (right). (E) Spatial distribution of chondrocyte cell type from the published cell segmentation analysis (left), compared with pixel-level inference by FICTURE (right), distinguishing ossification activity (blue) and other chondrocytes (yellow). The numbers in (D) and (E) indicate the number of cells or pixels found in the regions marked by the yellow dotted rectangles where the corresponding cell types are expected to be absent. The regions marked by the pink (D) or green (E) rectangles are magnified at the bottom. The full color codes and marker genes of each factor are shown in Supplementary Table 1.

We found that the factors identified by FICTURE tend to have specific spatial localizations that are more biologically sound than the published clusters based on cell segmentation. For instance, both methods identified the meninges, the membrane covering the brain and spinal cord, as one of their clusters or factors. The published cell segmentation analysis identified many cells outside the expected meningeal regions as meningeal cells, including 105 cells in the limb, but FICTURE classified zero pixels in meninges factor outside the central nervous system (Fig. 4D). Similarly, the original nuclear segmentation-based analysis mis-annotated subset of cells as chondrocytes in kidney, pancreas, and even brain, whereas the corresponding FICTURE factor representing cartilage activity was specifically localized to the developing skeletal elements (Fig. 4E). In addition, FICTURE correctly distinguished ossifying regions such as the palate and periosteum into a distinct factor representing ossification activity (Fig. 4E, right panel, blue), separated from other chondrocytes (yellow).

### FICTURE reveals fine-scale tissue architecture and cellular heterogeneity

In the same mouse embryo data, we also found that FICTURE revealed fine-scale tissue architecture and cell type heterogeneity obscured in the cell segmentation-based analysis. For example, the submandibular gland is composed of multicellular acini with high nuclear density as shown in the DAPI image (Fig. 4C, upper left). The nuclear segmentation analysis identified a mixture of epithelial cells, fibroblasts, and myoblasts, but the overall structure of the acini appears less clear than in the DAPI image (Fig. 4C, lower left). On the other hand, our pixel-level factor map distinguishes the structure of acinar cells and surrounding cells; the visualization a single epithelial factor clearly shows the structure of the acini (Fig. 4C, upper right), which is difficult to comprehend in segmentation-based results (Fig. 4C, lower left).

Moreover, in developing skeletal muscle, where cell-segmentation analysis labels almost all cells as myoblasts (Fig. 4B, upper two panels), FICTURE identified two distinct populations consisting of undifferentiated myoblasts, enriched for *Myf5*, *Pax3*, and *Pax7*, and differentiating myofibers, enriched for *Ttn* and *Dmd* (Fig. 4B, bottom panel; red and yellow respectively). The spatial pattern of myofiber factors inferred by FICTURE shows a remarkably elongated morphology, consistent with their transcriptome phenotype (Fig. 4B, lower two panels). On the other hand, DAPI-based segmentation fails to capture the structure of myofibers due to both their multinucleate nature and elongated shapes (Fig. 4B, second from the top). These results illustrate that FICTURE can outperform histology-based cell segmentation methods—currently considered the gold standard—by unbiasedly elucidating the details with high spatial precision.

### Unbiased single molecule analysis of *in situ*-based ST datasets without segmentation

We also applied our method to the public datasets from two *in situ* technologies: 10X Xenium based on *in situ* sequencing and Vizgen MERSCOPE based on sequential *in situ* hybridization. Since these datasets are accompanied with high quality images of DAPI and/or cell boundary staining, previous analyses collapsed individual transcripts by image-based cell segmentation. Data from these *in situ* platforms differ from the NGS-based technologies in that only a few hundred selected genes are measured, and each transcript has a unique spatial location. Therefore, in these datasets, FICTURE’s pixel-level inference has single molecule resolution. FICTURE successfully identified factors corresponding to cell types identified from previous segmentation-based analyses. Furthermore, as detailed below, FICTURE’s pixel-level analysis identified biologically important cell types and tissue structures that are not clearly captured by imaging-based cell segmentation.

### FICTURE recovers fibroblast and adipocyte morphology in inflamed tumor tissue

Using the 10X Xenium breast cancer dataset, we compared the performance of cell segmentation and FICTURE analyses. Compared to segmentation-based analysis^12^, FICTURE’s pixel level analysis more comprehensively characterized the entire section including fibrotic and fatty areas where cell segmentation is challenging. For example, the H&E near a ductal carcinoma *in situ* (DCIS) region in the lower center (red boxes in Fig. 5A, 5B) suggests the presence of lipid-laden adipocytes (Fig. 5C, center). However, the published cell segmentation analysis failed to detect the presence of adipocytes in this region: 45.3% of transcripts in the region were not assigned to any cell; when assigned, they were only classified as stromal cells (Fig. 5C, left), which are just one of the major constituents of stromal vascular fraction (SVF)^30^. Since adipocyte nuclei occupy only a tiny fraction of cell volume (0.1% if nuclear diameter is 1/10 of cell diameter) and non-lipid cell mass is more substantially contributed by SVF-associated cells, DAPI-based cell segmentation has poor sensitivity to capture transcriptome from differentiated lipid-laden adipocytes (Fig. 5C, right). Therefore, even though Xenium data contains adipocyte information^12^, segmentaion methods failed to isolate distinct adipocyte clusters. In contrast, FICTURE was able to confidently identify a factor specific to differentiated adipocytes enriched for *ADH1B*, *ADIPOQ*, *LPL*, and *LEP*. The lipid-laden morphology of differentiated adipocytes was also faithfully represented (Fig. 5D, center). In addition, FICTURE also identified all known SVF-associated cell types including endothelial cells, stromal cells, and macrophages (Fig. 5D, right), as well as other cell types (Fig. 5D, left), while preserving their histological structures around the vascular and fibrotic area. Such microscopic structures are inaccessible to traditional methods based on image-based cell segmentation (Fig 5C, left).

**Figure 5.**
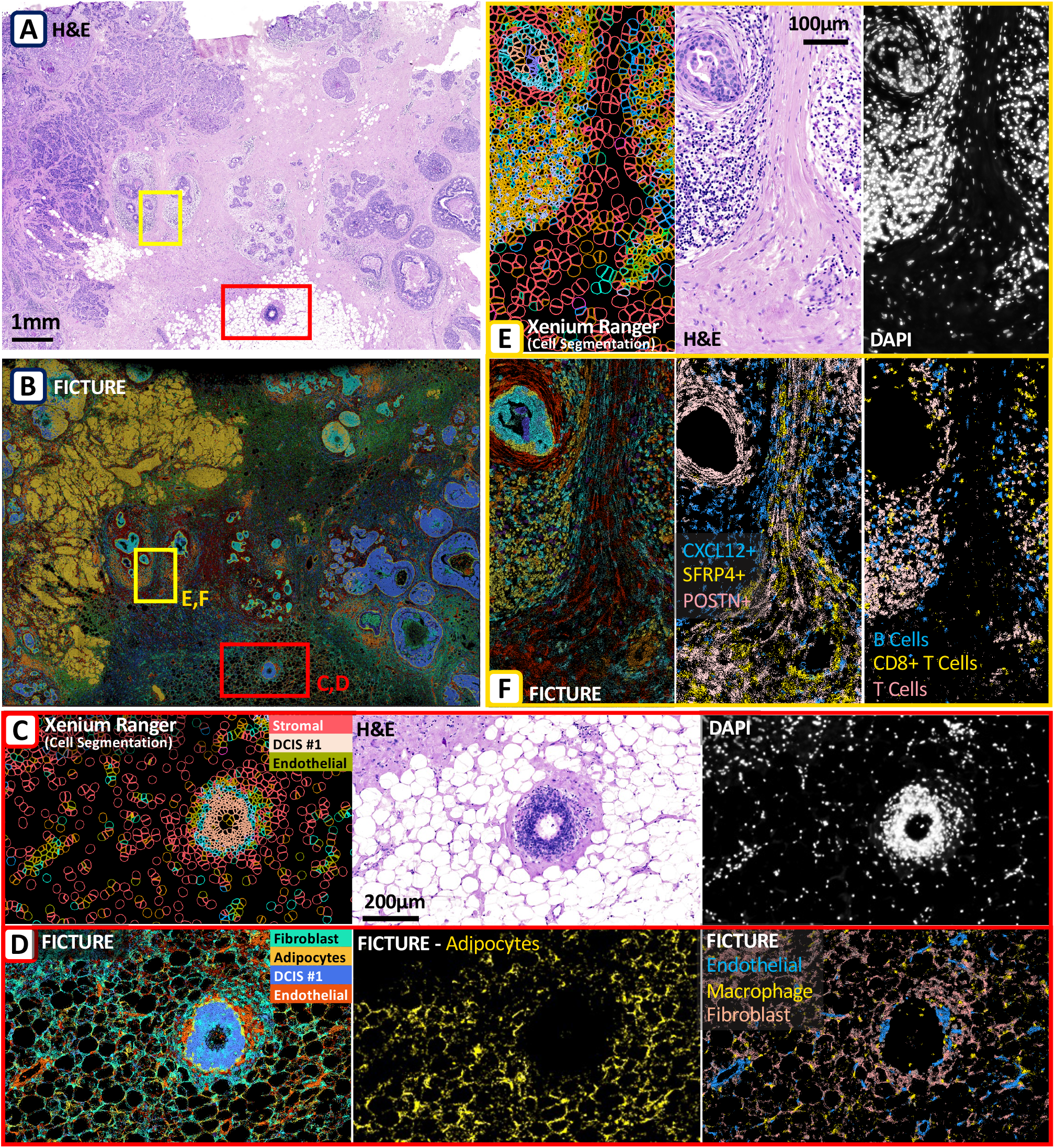
Application of FICTURE to a published Human Breast Cancer Dataset by 10X Xenium. (A) H&E image of the whole tissue section profiled by 10X Xenium. (B) The pixel-level factor inference using fully unsupervised FICTURE. (C, D) Adipose tissue surrounding ductal carcinoma *in situ* (red boxes in (A) and (B)) was magnified. (C) Cell segmentation-based clustering, H&E, and DAPI image (from left to right) from the published 10X Xenium analysis using Xenium Ranger. (D) Inference from unsupervised FICTURE. Left: all factors. Center: adipocyte-representing factor. Right: factors representing endothelial cells (blue), macrophages (yellow), and fibroblasts (pink). (E, F) Region between lesions of atypical ductal hyperplasia (yellow boxes in (A) and (B)) was magnified. (E) Cell segmentation-based clustering, H&E, and DAPI image (from left to right) from the published 10X Xenium analysis using Xenium Ranger. (F) Inference from unsupervised FICTURE. Left: all factors. Center: Factors representing three stromal cell populations, with *CXCL12* (blue), *SFRP4* (yellow), and *POSTN* (pink) as the top markers. Right: Factors representing immune cell types including B cells (blue), CD8+ T cells (yellow), and the other T cells (pink). The full color codes and marker genes of each factor are shown in Supplementary Table 1.

Fibrotic tissues are another tissue type where cell segmentation is challenging due to the irregularly elongated morphology of fibroblasts and their dense aggregation with other cell types such as immune cells. The region between the lesions of atypical ductal hyperplasia (yellow boxes in Fig. 5A, 5B) is mostly occupied by extracellular matrix produced by elongated fibroblasts in the tissue (Fig. 5E, center), which interferes with cell segmentation. In the cell segmentation analysis (Fig. 5E, left), many RNAs cannot be assigned to a nearby nucleus (Fig. 5E, right) and therefore dropped from the analysis (Fig. 5E). In contrast, our segmentation-free method preserves the tissue texture in this region (Fig. 5F, left) consistent with histology (Fig. 5, center); it detects and distinguishes three major stromal cell populations with distinct spatial distributions identified by the marker genes *POSTN*, *CXCL12*, and *SFRP4,* respectively (Fig. 5F, center). Among these, *CXCL12*-expressing fibroblasts are intermingled with different lymphocyte populations (Fig. 5F, right), consistent with their role in immune regulation. Genes enriched in adipocytes (*LEP, ADIPOQ, ADH1B, LPL*), stromal cells (*MUC6, CSF3, MPO, MMP2*) and myoepithelial cells (*UCP1, CXCL5, SDC4*) tend to have higher proportion of transcripts dropped during image-based segmentation (Supplementary Fig. 6B). Our analyses demonstrate that these ‘stray’ transcripts can be effectively allocated into relevant spatial factors using FICTURE. This functionality is particularly crucial when profiling complex regions like fibrotic and lipid-rich tissues, often linked with a range of pathologies.

### FICTURE reveals details of hepatic vasculature around the portal triad

We also applied FICTURE to the Vizgen MERSCOPE mouse liver datasets containing 419 million transcripts over 395,000 segmented cells within a 300 mm^2^ region. Approximately 93% of transcripts originated from large (20-30 µm) hepatocytes and the remaining transcripts are mostly from smaller non-parenchymal cell types, including Kupffer cells, endothelial cells, and hepatic stellate cells (HSC), which often have irregular shapes. Because the cell segmentation tends to be more difficult in smaller or irregularly shaped cells, we observed that the cell segmentation tend to be less accurate in non-parenchymal cells. For example, most (16/20) of genes with the highest proportion (>25%) of transcripts dropped from cell segmentation are enriched in fibroblasts, cholangiocytes, or arteries (Supplementary Fig. 6C). Across the entire liver section, FICTURE clearly visualized hepatocellular transcriptome heterogeneity aligned with metabolic zonation across portal-central axis (Fig. 6A, B). FICTURE also identified the precise locations of non-parenchymal cells, such as endothelial cells, macrophages, fibroblasts, and HSCs, and depicted their heterogeneity as well (Fig. 6C). The locations of these non-parenchymal cells were consistent with their histological niche near hepatic sinusoids and arteries (Fig. 6C, D).

**Figure 6.**
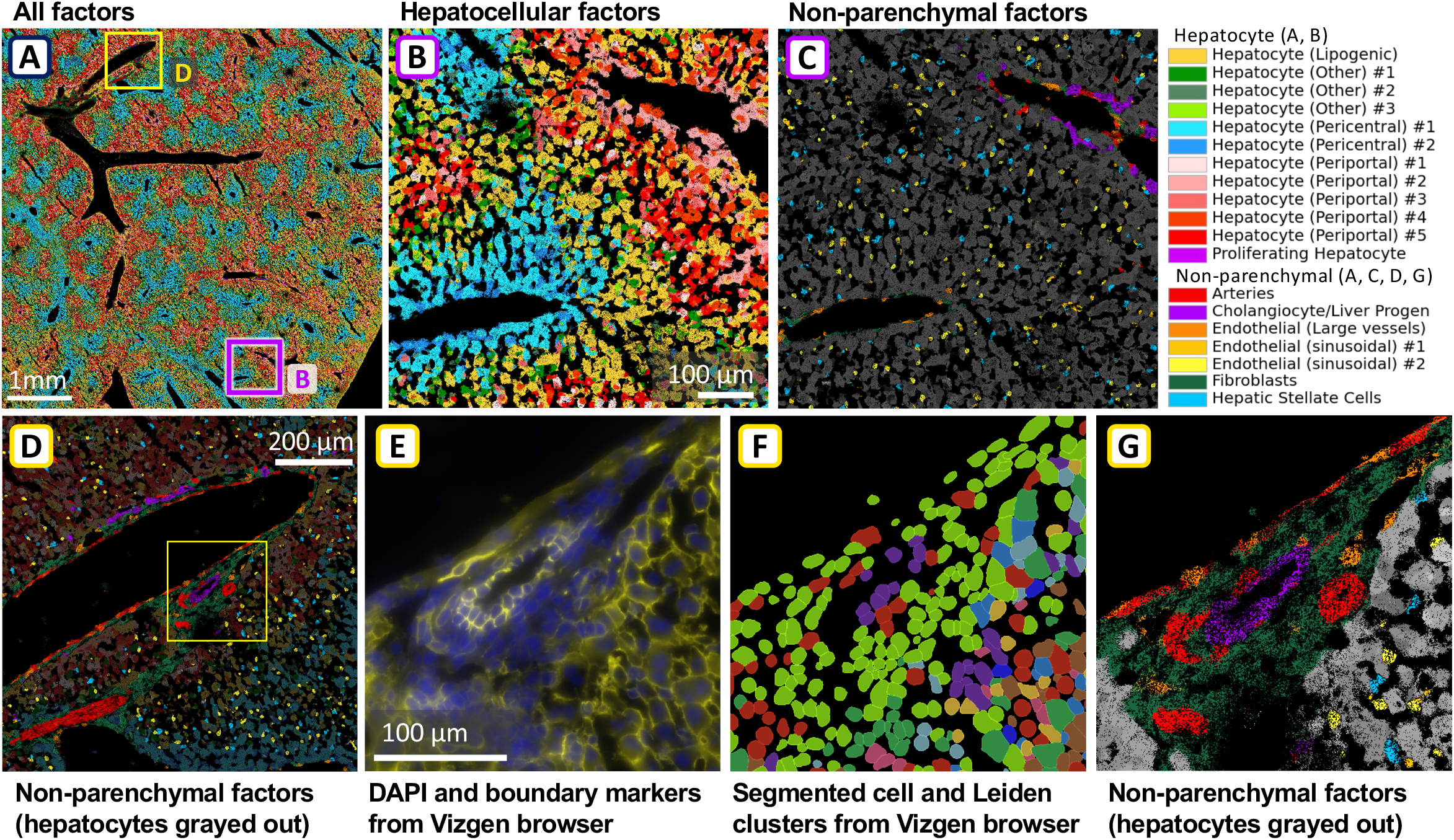
Application of FICTURE to a Mouse Liver dataset by Vizgen MERSCOPE. (A) A wide view of the pixel-level inference result of the mouse liver section, showing all factors as different colors. The two squares indicate the region magnified in the other panels. (B) A close-up view of pixel-level factorization near a hepatic sinusoid focusing on periportal and pericentral hepatocytes. (C) Corresponding view of the same region as in (B), focusing on non-parenchymal cells. (D) Close-up view of non-parenchymal factors around portal vein. Yellow box was further magnified in (E-F). (E) A further magnified view of nuclei (blue) and cell boundary markers (yellow) near the portal vein from the Vizgen browser. (F) The magnified view of the same region as in (E) showing the segmented cells colored by their Leiden cluster assignments from the Vizgen browser. (G) Factors from FICTURE in the same regions as in (E, F), focusing on non-parenchymal cells. The full color codes and marker genes of each factor are shown in Supplementary Table 1.

In the portal triad area, where portal vein, arteries and bile ducts, as well as fibrotic and inflammatory tissues are aggregated together, densely populated nucleus and extremely stretched ductal and endothelial cell morphology makes cell boundary estimation from nuclear and membrane markers extremely challenging and often not feasible (Fig. 6E, F). Correspondingly, image-based cell segmentation is incomplete and misses many cells constituting important structures such as cholangiocytes forming bile duct (Fig. 6F). FICTURE, however, recovers all these cells and the surrounding fibroblasts with clear structures forming portal vein endothelial cells, bile ducts and arteries with their characteristic histological morphology (Fig. 6D, G). These results exemplify the unique utility of FICTURE in labeling histology with relevant transcriptome information.

### Scalability and accuracy of FICTURE in large real-world datasets

Even though FICTURE comprehensively analyzed all these massive datasets without difficulties, other former methods we used for initial comparison, Baysor and GraphST, failed to analyze them in our attempt. To assess the computational feasibility of applying these existing methods for the datasets analyzed above, we projected the computational cost to run Baysor and GraphST using the values from the simulation study and compared it with the actual computational cost in FICTURE (Fig. 3F). Even if we limited the number of genes to 500 for Baysor and GraphST, both methods would require prohibitive memory, over 2TB for two of the three datasets, while FICTURE consumes less than 10GB. The computational time to analyze these datasets was projected to be a day or less for FICTURE and GraphST, while Baysor was projected to take up to decades.

To compare the accuracy in the real datasets within a reasonable computational time, we selected a very tiny (0.46mm^2^) subset of the Vizgen’s mouse liver data containing both central and portal veins presented above and compared FICTURE, Baysor, and Graph ST. All methods successfully distinguished between pericentral and periportal hepatocytes, but only Baysor and FICTURE identified the precise locations of small and irregularly-shaped non-parenchymal cells (arrows in Supplementary Fig. 7) and the inferences from the two methods are in high concordance. GraphST’s results showed artificial spatial patches with resolution much coarser resolution than single cells, even though it operates at single cell level.

## Discussion

Spatial transcriptomics technologies have evolved to profile the transcriptomes of large tissue sections at submicron resolution. However, submicron ST technologies do not automatically enable submicron resolution analysis. The limited number of RNA molecules observed per μm^2^ leads to a tradeoff between spatial resolution and information content. At the same time, the increasing number of uniquely identified spatial locations (pixels) poses a challenge to computational scalability. To realize the full potential of emerging submicron ST technologies, we have developed FICTURE, a highly scalable method for spatial factor analysis at submicron resolution without relying on cell segmentation what is compatible with both imaging-based and sequencing-based submicron ST technologies. FICTURE adaptively combine sparse information from individual pixels and probabilistically assigns individual pixels to latent factors that may represent expression patterns of cell types, cell states, biological processes, or disease conditions.

To date, most analyses of high-resolution spatial transcriptomics data rely on either fixed-size gridding or cell segmentation^8,9,12^. These approaches sacrifice spatial resolution to increase the number of transcripts per unit of analysis and reduce computational burden. We demonstrated in our Seq-scope colon dataset that gridding-based analysis risks obscuring biological interpretation especially in regions where multiple cell types intermingle. Cell segmentation analysis relies on the availability and accuracy of external nuclear and/or cell boundary staining. Cell segmentation based solely on nuclear staining is the most common but requires strong assumptions about cell size and shape. We showed in the 10X Xenium breast cancer data that nuclear staining is inadequate for cells with irregular morphology, such as fibroblasts in tumor stroma and adipocytes in lipid-laden area. This limitation is significant because both cell types are integral to understanding tumor progression: adipocytes interact with breast cancer cells through signaling factors^31^ and cancer-associated fibroblasts in the stroma have multiple functions in supporting tumor growth^32^. Fibrosis and fat accumulation are also commonly associated with inflammation, metabolic dysfunction and other tissue pathologies; therefore, precisely analyzing these cell types is important. Even when high-quality cell boundary markers are available, as in the MERSCOPE liver dataset, we observe that cell segmentation can fail in regions of densely packed cells and systematically misses certain cell types. Our segmentation-free approach does not require assigning transcripts to cells before making inferences in the complex regions of a tissue. As a result, FICTURE provides an unbiased and comprehensive interpretation of submicron ST data and can be used as an alternative or complementary approach to segmentation-based analysis.

Even in cases where cell segmentation works well, pixel level inference can provide additional information about ongoing biological processes. For example, in Xenium human lung preview dataset non-diseased tissue section, where cell segmentation captures 88.3% of transcripts, the cell level clustering analysis and FICTURE’s pixel-level inference were largely consistent. The most noticeable difference is that FICTURE identified a factor enriched for inflammatory markers specifically localized to the area of peribronchial immune cell infiltration. The cell level analysis in the Xenium does not have a corresponding localized cluster (Supplementary Fig. 8A-B). Upon detailed examination of these peribronchial immune cell infiltrates, the presence of *CD8A, IL7R (CD127), CD28, and CD27* suggests an enrichment of effector memory T cells, and one of the most region-specific genes, *GZMK*, suggests the presence of a specific inflammatory T cell subtype in these regions (Supplementary Fig. 8D-F). Cell segmentation profiles such regions as a mixture of individual cell types, such as CD8 T cells, B cells, monocytes, and lymphocytes, regardless of the inflammatory signature (Supplementary Fig 8C). These results suggest that FICTURE can be used as a complementary method to cell segmentation when interpreting submicron ST data, as the unsupervised FICTURE factors may recapitulate the unique tissue microenvironment that are not well captured in single cell level analysis.

FICTURE is the first pixel-level analysis tool that scales to sequencing-based technologies such as Seq-scope and Stereo-seq. Sequencing-based technologies unbiasedly profile the whole transcriptome while *in situ* hybridization-based technologies typically assay only a few hundreds of pre-selected genes in each sample. The large number of genes makes existing methods designed for *in situ*-based technologies^22,33^ infeasible for whole transcriptome datasets due to computational scalability. Baysor, the most accurate segmentation-free method to our knowledge, recovers cell and cell type boundaries in both real and simulated data but requires prohibitive resources to process large regions and is not applicable to data from sequencing-based technologies with tens of thousands of genes. FICTURE’s pixel-level inference has comparable or better accuracy than Baysor but is orders of magnitude more efficient. FICTURE’s inference is parallelized and scales linearly with the number of pixels. Furthermore, FICTURE adopts a stochastic approach so that it can run with a fixed memory budget, a property uncommon among existing methods but crucial for analyzing data with large fields of view as exemplified in the currently presented Stereo-seq and MERSCOPE cases. FICTURE is also more robust to over-specification of the number of factors, reducing the hyperparameter tuning in practice where we rarely know the best level of model complexity. These features will become increasingly useful in future datasets as the information content per experiment continues to grow ^22,33^.

The computation efficiency does require a few simplified assumptions. First, the LDA model that FICTURE builds upon is limited by its assumption that factors are independent. Modeling correlation and hierarchy between factors could be a future direction towards more biologically meaningful models. One could also conceive incorporating prior information such as known marker genes, penalizing redundant or highly similar factors, and encouraging sparsity in factor-level expression profiles. Second, FICTURE decoupled the factorization procedure from the pixel-level inference to allow flexible pixel-level inference with or without external reference. When the transcript density is high it might be possible to improve the factorization during the pixel-level inference although we did not observe the benefit in our data.

FICTURE requires users to specify the density of anchors and the size of the overlapping anchor neighborhoods. The anchor density is intended to guarantee, on average, at least one anchor near each unknown cell center; the neighborhood size is intended to contain enough molecules to infer the latent factors and it controls spatial sparsity thus computational cost. While these hyperparameters can be chosen intuitively based on the data, if either the molecule density or the anchor density is too low, FICTURE tends to oversmooth by sharing information over a large area. Oversmoothing, or the bias towards assigning the same factors to neighboring pixels, is due to overconfidence in pixel-level factor assignments. This overconfidence results from the well-documented tendency of variational inference to underestimate the variance of the posterior density^34^. A future improvement is to implement a diagnostic mechanism to identify pixels where the factor assignment is truly ambiguous, and to avoid assigning such pixels to the dominant factor in their neighborhood which creates the artificially smooth spatial pattern. Oversmoothing is also a common risk in low-resolution spatial methods^9^, including the popular graph neural network^35^. As we have shown in all the datasets we analyzed, cellular heterogeneity in real tissues is very high. Consequently, assuming spatial autocorrelation at the cell level is often too strong of an assumption. Therefore, we propose to use spatially agnostic models on collapsed grid-level expressions for factorization prior to pixel-level analysis.

With the rapid development of high-resolution, high-throughput spatial transcriptomics technologies, we hope that FICTURE will facilitate the development of robust and efficient analysis workflows for translating raw data into biological insights. FICTURE’s segmentation-free approach may be particularly suited for studying complex tissues, extracellular matrix, or cell-cell communication. Its scalability, robustness, and intuitive hyperparameter selection make it easy to apply to new protocols and tissue types.

## Methods

### FICTURE Model

We assume that a ST dataset consists of *M* genes measured at *N* pixels, represented by a sparse count matrix *X* ∈ *Z^N^*^×*M*^. Each pixel is associated with a unique two-dimensional coordinate *y_i_* ∈ ℝ^2^, *i* ∈ [*M*]. Each pixel may contain multiple molecules from different genes. We define a hexagonal lattice over the entire tissue section and lattice points inside tissue regions as anchors, where the distance between adjacent lattice points, a, controls anchor density. These n anchor points are surrogates for unknown cell centers and serve the purpose of sharing pixel level information locally. Each anchor j ∈ [n]’s territory is upper bounded by a circle centered at its location 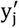 with a fixed radius *d_max_*, and nearby anchors’ territories overlap. The tradeoff between spatial resolution, factorization accuracy, and computational cost depends on the anchor density a and the radius *d_max_*. We choose *a* so that each cell covers more than one anchor with high probability and choose *d_max_* be larger than *a* and smaller than the diameters of cells that could be present in the data.

Our model (Supplementary Fig. 1A) extends LDA by allowing each pixel to choose (within *d_max_*) which anchors it belongs to probabilistically, thus in turn coupling cell type estimates of nearby anchors through the pixels they may share. Let the set of nearby candidate anchors for pixel *i* be *n*(*i*) and the set of pixels within an anchor *j*’s territory be *N*(*j*).

Pixel *i*’s probability of belonging to anchor *j* is *w_ij_* ∈ [0,1] a priori, here we choose 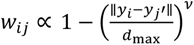, a concave monotone decreasing function of the distance between the two points with a fixed hyper-parameter *ν*. This extra uncertainty is highly localized, designed to refine cell type boundary to beyond the resolution achieved by sliding hexagons while avoiding unnecessary smooth assumption about the underlying cell type distribution.

The generative process is as follows:

1. Draw factor level expression distribution **β**_1_ ~ Dirichlet(**η**) for *k* ∈ [*K*]
2. Draw anchor level factor proportions **θ***_j_* ∈ Dirichlet(**α**) for *j* ∈ [*n*]
3. For each pixel *i* ∈ [*N*]:

a. Draw anchor assignment *C_i_* ~ Categorical(**w***_i_*) (where **w***_i_* = (*w_ij_*), *j* ∈ *n*(*i*))
b. Draw factor assignment 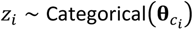, *Z_i_* ∈ [*K*]
c. Draw gene expression 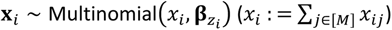

Each pixel is equipped with two categorical variables, the anchor assignment *C_i_* ∈ [*n*(*i*)] and factor assignment *Z_i_* ∈ [*K*]. When there is no ambiguity, we use *Z_i_*_1_ both for the event *Z_i_* = *k* and the indicator variable *Z_i_*_1_ = *I*{*Z_i_* = *k*} (similar for *C_ij_*). The joint likelihood can be written as follows:

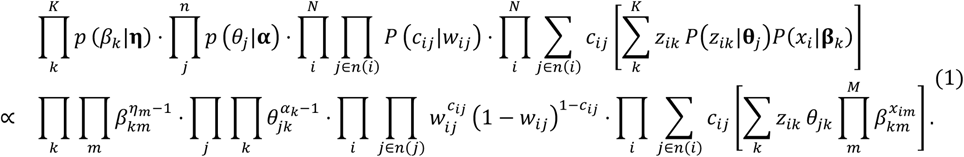

### Variational inference in FICTURE

We use stochastic variational inference^36,37^ to approximate the posterior distributions. Global variables are factor level expression profiles **β**_1_ ~ *Dir*(**λ**_1_). Local variables include anchor level factor proportion **θ***_j_* ~ *Dir*(**γ***_j_*), pixel level anchor assignment *C_i_* ~ *Cat*(**ψ***_i_*), and pixel level factor assignment *Z_i_* ~ *Cat*(**ϕ***_i_*). Spatial coupling introduces dependence between anchor points, so we construct minibatches as sliding windows of size ≫ *d*_max_ that jointly cover the dataset. For each minibatch, we only update variables for the “interior” anchors and pixels that are independent from the rest of data conditional on this minibatch. We randomize the order of minibatches so that different cell types are similarly represented across the course of stochastic optimization. Coordinate ascent update rules based on the above mean-field approximation resemble that for LDA (Supplementary Text).

In the fully unsupervised FICTURE, we initialize this model from the standard LDA. We collapse pixels into hexagons of side length *r* and train a LDA model as if the hexagons were independent. We use the posterior distributions 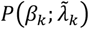 of the *β*_1_’s in this spatially agnostic model as the prior *P*(*β*_1_; *η*_1_) in the full model.

### Pixel level decoding from reference cell types

When external reference is available or when other clustering methods have been applied to collapsed low resolution data, we aggregate the gene counts within each cell type or cluster to a pseudobulk expression count matrix 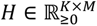, and substitute it for the posterior Dirichlet parameter of factor level gene expression profiles. We then either fix this global parameter and only update local parameters as in Equation (6) in the Supplementary Text, or treat it as a prior and replace Equation (7) in the Supplementary Text with 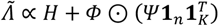.

### Simulation

#### Simulating cells

We first simulate cell centers as uniformly random perturbation of regular grid points. We set the distance between adjacent grid points as *A* (10 *or* 15μ*m*), and perturb each grid point as *x* = *x*_0_ + δ*_x_*, *y* = *y*_0_ + δ*_y_*, where δ*_x_*, δ*_y_*~ Unif(−0.5*A*, 0.5*A*). We simulate cells of three shapes, circle, rod, and rhombus, and randomly assign 40% cells as circles, 30% as rod, and 30% as rhombus. We set the radius of the circles to 6μ*m*, the length and width of the rod to be 16 and 4μ*m*, and the side length and the acute angle of the rhombus to be 15μ*m* and *π*/16. We also simulate a buffer region of width 2μ*m* around each cell to mimic RNA diffusion, and randomly sample 50% of molecules in the buffer region to be from the focal cell. We place cells according to a randomized sequence and cells positioned later partially superimpose cells positioned earlier when they overlap.

We simulate molecules uniformly distributed in the region covered by cells with a desired molecule density, often between 1 to 8 molecules per μ*m*^2^ according to different technologies. Regions outside any cell’s buffer region have molecule density 20% that of cellular regions.

#### Simulating gene expression

We simulate gene expression based on annotated scRNA-seq data from Tabula Muris^27^. To benchmark with Baysor^22^, which is developed for imaging-based technologies and does not scale to large number of genes, we choose 500 highly variable genes from the whole transcriptome based on the genes’ variance to mean ratios across cells. We choose 10 cell types, assign 8 of them to specific spatial domains and scatter cells from the other 2 cell types uniformly across the space (Fig. 3A). For these 8 localized cell types we simulate two types of cell type boundaries, deterministic and probabilistic. We construct the deterministic boundaries as smooth lines and assign each cell to a cell type domain by its centroid location. We construct the fuzzy, probabilistic boundaries by sampling cell type identities from a mixture distribution with two components. The mixing probabilities smoothly change from (0.5,0.5) to (0,1) and (1,0) moving away from the hypothetical boundary line.

We assign intracellular molecules the cell type of the cells they originate from and assign extracellular molecules the cell type of their nearest cell. We then simulate gene identity of each molecule from a categorical distribution defined by the relative expression level of its cell type. Equivalently, each cell’s gene expression follows a multinomial distribution.

### Submicron resolution spatial transcriptomics datasets across four platforms

The colon samples were obtained from C57BL/6 mice after 24 hrs of biopsy-induced physical wounding, as previously described^25^. The Seq-Scope procedure was performed on the injured colon sample as previously described^8^ with minor modifications involving the use of the Illumina HiSeq 2500 flow cells and chemical strategies to increase RNA sequence capture^38^. Seurat factors were learned from a dataset with hexagonal grids with 14 µm sides, using Seurat v4. In brief, mitochondrial genes and hypothetical gene models were removed from the dataset, and feature cutoff was applied at 200. Data were normalized by *SCTransform* function, and clustering was performed using *FindClusters* function (resolution = 2). A macrophage cluster was subjected to an additional round of clustering to get separation between different macrophage subtypes. Top markers from each cluster were used to infer and annotate cell types.

The Stereo-seq pixel-level expression data was downloaded from the “Bin1 matrix” section in the STOmicsDB^39^ (https://db.cngb.org/stomics/mosta/download/) for the embryo E16.5_E1S3. “Cell_bin” matrix was also downloaded to obtain the raw expression data used for cell-level segmentation. The published annotations of segmented cells were obtained from the “Embryo Data” section (<u>E16.5_E1S3_cell_bin.h5ad</u>).

The 10X Xenium breast cancer dataset was downloaded from the 10X Genomics web page (https://www.10xgenomics.com/resources/datasets/ffpe-human-breast-with-pre-designed-panel-1-standard). Transcript-level data (transcripts.csv.gz) was used to identify the spatial coordinates of each transcript. The published clusters were obtained from the analysis output based on graph clustering (analysis/clustering/gene_expression_graphclust/clusters.csv). The same procedure was repeated for the 10X Xenium healthy lung preview dataset available at (https://www.10xgenomics.com/resources/datasets/xenium-human-lung-preview-data-1-standard).

The Vizgen MERSCOPE mouse liver data was download from https://info.vizgen.com/mouse-liver-access. The sample L1R1 was used for the analysis. The transcript level data (detected_transcripts.csv.gz) was used as input for FICTURE in addition to the DAPI staining image at a selected z-coordinate (z3). The Mouse Liver map browser on the Vizgen web site was used to show the cell segmentation and clusters as such dataset was not available in the released full dataset.

Images obtained from grid, cell segmentation and FICTURE analyses of these datasets were linearly adjusted to visualize dim regions and highlight different factors/clusters.

### Processing submicron resolution spatial transcriptomic datasets

#### Filtering

We only performed filtering for Seq-scope data. We first filtered out low density region assuming tissue regions have much higher transcript density than low quality regions dominated by noise. We aggregated transcripts by a coarse sliding grid and calculate transcript density in each grid cell, then fit a two-component Gaussian mixture model on the log-transformed grid densities. We kept grids assigned to the high-density component. We then filtered out pixels outside manually drawn tissue boundaries. For Stereo-seq and Vizgen MERSCOPE we performed analysis on all transcripts in the released data. For 10X Xenium we kept transcripts with quality score above 15.

#### Grid level analysis

We performed model training and anchor initialization at hexagon level. We aggregated transcripts to hexagons with side length 14 µm (Seq-scope colon data), 8.7 µm (stereo-seq embryo data) or 7 µm (all in situ datasets) according to the regular hexagonal grid, and trained LDA on these hexagons. For Seq-scope colon data we transformed the hexagonal data with sctransform^40^ and applied the Louvain algorithm as implemented in Seurat^16^. We chose anchors as the lattice points on a denser hexagonal grid where the distance between two adjacent lattice points is 4 µm. We initialize the factor mixture probabilities at each anchor by applying the trained LDA model on the hexagon centered at the anchor, where the size of the hexagon is the same as that used in the model training. We then flatten the mixture probabilities by lower-bounding the minimum to 1/2K (K is the number of factors) then normalize again to weaken the prior information from this hexagonal level analysis.

### Comparison with other methods

We ran Baysor and GraphST with default settings on CPUs with a single thread. For Baysor we set the key parameter, expected cell radius, to the simulated cell radius. In the Vizgen mouse liver subset, we set the expected cell radius to 7 µm. For GraphST we chose Leiden for final cluster assignment because it works most reliably. We did not use the “refinement” option in GraphST because it tended to create smoother spatial pattern that does not match our simulation setting resulting in lower accuracy. Baysor and FICTURE’s runtime is stable across runs, but GraphST’s runtime was highly variable. We ran GraphST 10 times and took the average runtime and memory as shown in Fig. 3E.

## Data availability

The source code and python package for FICTURE method is publicly available in the GitHub repository at https://github.com/seqscope/ficture. The full results from the 5 real datasets are available in Zenodo with DOI https://doi.org/10.5281/zenodo.10070621. The Seq-Scope mouse colon data will be publicly available in the Gene Expression Omnibus. Other datasets can be accessed from the sources indicated in the Methods section.

## Acknowledgements

The work was supported by the Taubman Institute (to HMK and JHL) and the NIH (R01HG011031, and HHSN268201800002I to HMK, UH3CA268091 and R01AG079163 to JHL).

## Ethics declarations

HMK owns stock for Regeneron Pharmaceuticals. JHL is an inventor on a patent and pending patent applications related to Seq-Scope.

