## Supplementary Text and Figures for "*FICTURE:* Scalable segmentation-free analysis of submicron resolution spatial transcriptomics"

### Inventory of Supporting Information

#### Supplementary Figure Legends

**Supplementary Figure 1 – Generative Models used in FICTURE**. (A) The generative model for pixel-level inference used in FICTURE. Shaded circles represent observed data. $(x_j, x_i)$ represent spatial locations of anchors and pixels; $\widetilde{y_\iota}$ represents gene counts for pixel $i$. The black-outlined circles $(\eta, \alpha)$ represent hyperparameters for the Dirichlet priors. The orange-outlined circles $(\beta_k)$ represent factor level expression distribution and the green-outlined circles $(\theta_j)$ represent the anchor level factor proportions. The blue-outlined circles $(z_i, c_i)$ represent latent factor and latent anchor assignments for each pixel. We provide the typical range of the number of factors (K), anchor points (n), and pixels (N) next to the corresponding boxes. (B) Latent Dirichlet Allocation model used in the fully unsupervised FICTURE. The standard LDA model is used to infer factors $(\beta_k)$ to model gene counts for each "spot" and the learned factors are used as input to part (A). Each "spot" level gene count $(\widetilde{y_{\iota,J}})$ is generated using a fixed-sized hexagonal grid. $\Theta_j$ represents the probabilistic distribution over K factors for spot j. $z_{ij}$ represents the latent factor of pixel $i$ in spot $j$.

**Supplementary Figure 2 – Overview of the Datasets Analyzed.** A tabular summary of the datasets analyzed in this manuscript, in order of appearance. Large datasets (>10mm$^2$) were analyzed with FICTURE only since they are not analyzable by other methods. Simulated datasets and smaller subset of real datasets were analyzed with multiple methods.

**Supplementary Figure 3 – Additional data of Seq-scope mouse colon and marker genes.** (A) H&E (top) and RNA density (bottom) of the full dataset including 9 tissue sections. (B) A dot plot visualizing the marker genes in the Seq-Scope mouse colon dataset. The x-axis represents the marker genes identified from Seurat, and the y-axis represent different clusters with their annotated cell types. The colors represent the fold enrichment of average expression levels of each gene compared to the rest of the cell types. The size of dot represents the relative expression of the marker gene among all genes in each cluster. (C) FICTURE's pixel-level decoding result of the whole data (top) and the magnified view in each section (bottom). The full color codes and marker genes of each factor are shown in Supplementary Table 1.

**Supplementary Figure 4 – Comparison between FICTURE, Baysor, and GraphST using the densely packed simulation model.** A visualization similar to Fig. 3B, but instead of simulating transcripts assuming sparse background outside cell boundaries, this simulation uses the uniform transcript density across the entire region. We refer to this simulation as "densely packed" while "sparse background" represents the simulation in Fig. 3B. (A-C): All pixels are visualized with colors representing their inferred factors from (A) FICTURE, (B) Baysor, and (C) Graph ST. (D-F): Pixels where the factors inferred from (D) FICTURE, (E) Baysor, and (F) GraphST differ from the ground truth are visualized with colors representing their true generating factors.

**Supplementary Figure 5 – Bias of DAPI-based cell segmentation in Stereo-seq dataset.** (A) DAPI staining images of Stereo-seq E16.5 (E1S3) mouse embryo. (B) Spatial distribution of erythrocyte from the published 25-cluster annotation based on cell segmentation. (C) Spatial distribution of erythrocyte factor from FICTURE. (D) A magnified view of DAPI image focusing on the heart and blood vessel. (E) A corresponding magnified view of published annotation. (F) A corresponding magnified view of FICTURE factors. (G) Proportion of transcripts assigned to cells within each tissue area. The 23 tissue areas are annotated based on the published grid-based segmentation. The full color codes and marker genes of each factor are shown in Supplementary Table 1.

**Supplementary Figure 6 – Proportion of dropped transcripts during cell segmentation.** Each gene is plotted based on the total number of transcripts in the whole dataset (x-axis) and the proportion of the transcripts excluded after the cell segmentation (y-axis). Each gene is colored by the tissue or factor that it is most enriched in. (A) Stereo-seq E16 mouse embryo data across 30,050 genes. The 23 tissue annotations are from the published grid-based segmentation. The top 40 most excluded genes (with >10,000 transcripts) are enriched in liver (*Hba-a2, Alas2, Hba-a1, Hbb-bs, Hbb-bt, Ube216, Mkrn1, Apex2, Trim71, Bpgm, Snca, Fam220a, Hbb-bh1*), heart (*Myl7,Tnni3, Myh6, Myl3, Myl2, Myh2, Pln, Tnnt2, Csrp3, Sh3bgr, Cox7a1, Ankrd1, Mybpc3*), blood vessel (*Hba-x, Hba-y*), epidermis (*Krt77*), and muscle (*Ckm, Cox8b, Tnni2, Myl4, Cox6a2, Tceal7, Acta1, Myh8, Mylpf, Synpo2l, Tnnt3*). (B) 10X Xenium human breast cancer data across 313 genes analyzed with 20 factors inferred from FICTURE. (C) Vizgen mouse liver data across 347 genes analyzed with 24 factors inferred from FICTURE. The factors in (B) and (C) are annotated based on marker genes and spatial distribution.

**Supplementary Figure 7 – Comparison of Methods on the Mouse Liver Dataset.** A small ($0.46mm^2$) subset of the Vizgen's mouse liver data containing both central and portal veins are selected to compare FICTURE with Baysor and GraphST. All methods are run with 12 factors/clusters. (A) Pixel-level output from FICTURE, (B) Pixel-level output from Baysor, and (C) Cell-level output from GraphST, rendered at pixel resolution based on cell segmentation. Magnified views of portal veins for (D) FICTURE, (E) Baysor, and (F) GraphST. Magnified views of central veins for (G) FICTURE, (H) Baysor, and (I) GraphST.

**Supplementary Figure 8 – Analysis of Human Lung Preview Dataset.** (A) Pixel-level visualization of inferred factor by FICTURE on a healthy human lung tissue assayed by 10X Xenium platform. An upper-left subsection was magnified in the inset. (B) Further magnified view of FICTURE's pixel-level inference on a peribronchial region showing a signature of immune infiltration (orange in the lower center). (C) Corresponding view in the 10X Xenium Explorer based on cell segmentation, each segmented cell colored by its assigned cluster. (D) DAPI image of the same region. (E) Gene-level visualization of the same region, focusing on *CD8A, CD27, IL7R*, and *CD28* in different colors. (F) Gene level visualization of *GZMK*. The full color codes and marker genes of each factor are shown in Supplementary Table 1.

#### Supplementary Table Legends

**Supplementary Table 1 – Detailed information of factors and clusters in the manuscript.** Each spreadsheet contains a table providing with detailed information of the factors inferred from FICTURE or clusters from another method. The columns include factor ID (0-based in teger), RGB color code used for visualization (RGB), the proportion of pixels assigned to each factor (weight), the total number of transcripts assigned to each factor (PostUMI), the top 20 genes based on the p-value from a simple chi-square test based on pseudobulk for each factor (TopGene_pval), the top 20 enriched genes based on the fold-changes from pseudobulk test (TopGene_fc), the top 20 genes that are most expressed within each factor (TopGene_weight), and the predicted cell type based on manual annotation. If the predicted cell type is not certain, it is left as blank.

**Supplementary Table 2 – Proportion of transcripts dropped during cell segmentation in each gene.** Each spreadsheet corresponds to one dataset, listing for each gene the total number of detected transcripts, the number of transcripts used in published cell segmentation (“Assigned”), and the proportion dropped during cell segmentation ("fDropped”). We also labeled each gene with the tissue (for Stereo-seq data, from published annotation) or factor (for Xenium & Vizgen data, from FICTURE) that it is most enriched in. The tissue/factor enrichment is quantified by Chi-squared statistics and fold change. For stereo-seq data we include only protein-coding genes with total count $\geq 100$.

### Supplementary Text

#### Coordinate ascent details

Each minibatch iterates over the updates of three blocks of local parameters ϕ, ψ, and γ until they converge:

$$\psi_{ij} \propto \text{expit}\left(\text{logit}\, w_{ij} + \sum_k \phi_{ik}\left(E[\log \theta_{jk}] + \sum_m x_{im} E[\log \beta_{km}]\right)\right) \quad (2)$$

$$= \text{expit}\left(\text{logit}\, w_{ij} + \sum_k \phi_{ik}\left(\tilde{\gamma}_{ik} + \sum_m x_{im}\, \tilde{\lambda}_{km}\right)\right)$$

$$\phi_{ik} \propto \exp\left(\sum_j \psi_{ij}\left(E_q[\log \theta_{jk}] + \sum_m x_{im}\, E_q[\log \beta_{km}]\right)\right) \quad (3)$$

$$= \exp\left(\sum_j \psi_{ij}\left(\widetilde{\gamma_{ij}} + \sum_m x_{im}\, \widetilde{\lambda_{km}}\right)\right)$$

$$\gamma_{jk} \propto \alpha_k + \sum_{i \in N(j)} \psi_{ij}\, \phi_{ik} \quad (4)$$

Here $\tilde{\gamma}$ and $\tilde{\lambda}$'s are expectations of the log of Dirichlet variables: $\widetilde{\gamma_j} \coloneqq E_q[\log \theta_j]$, $q(\theta_j) = Dir(\gamma_j)$; similarly, $\widetilde{\lambda_k} \coloneqq E_q[\log \beta_k]$, $q(\beta_k) = Dir(\lambda)$. Let $h(z)$ be the (element wise) digamma function (usually denoted as ψ, here we do not want to confuse it with the random variable ψ), then

$$E[\log \theta_j] = h(\gamma_j) - \sum_k h(\gamma_{jk}),\ E[\log \beta_k] = h(\lambda_j) - \sum_m h(\lambda_{km}).$$

We update the global parameters with learning rate $\rho_t$

$$\widetilde{\lambda_{km}^{(t)}} \propto \eta_m + \sum_i \sum_{j \in n(i)} \psi_{ij}\phi_{ik}x_{im} \qquad (5)$$

$$\lambda_{km}^{(t)} = \rho_t \lambda_{km}^{(t-1)} + (1-\rho_t)\widetilde{\lambda_{km}^{(t)}}$$

We can write the algorithm compactly in matrix form with the following notations:

- $X \in Z_{\geq 0}^{N\times M}$, (sparse) observed pixel level gene counts
- $W \in R_{\geq 0}^{N\times n}$, (sparse) prior pixel assignment based on spatial locations of pixels and anchor points.
- $\Lambda \in R_{\geq 0}^{K\times M}$: posterior Dirichlet parameters for factor level gene expression $\boldsymbol{\beta_k}$
- $\widetilde{\Lambda} = h(\Lambda) - h(\Lambda) * 1 * 1^T \in R_{\geq 0}^{K\times M} : \widetilde{\Lambda_{k\cdot}} \propto E_{\beta_k \sim Dir(\Lambda_{k\cdot})}[\log \boldsymbol{\beta_k}]$ (normalized expected $\log \beta$ under variational distribution)
- $\Gamma \in R_{\geq 0}^{n\times K}$: posterior Dirichlet parameters for anchor level factor assignment $\boldsymbol{\theta_j}$
- $\tilde{\Gamma} \coloneqq h(\Gamma) - h(\Gamma) * 1 * 1^T \in R^{n\times K}$: $\widetilde{\Gamma_{j\cdot}} = E_{\theta_j \sim Dir(\Gamma_{j\cdot})}[\log \boldsymbol{\theta_j}]$ (normalized expected $\log \theta$ under variational distribution)
- $\Phi \in [0,1]^{N\times K}$: inferred pixel level factor assignment
- $\Psi \in [0,1]^{N\times n}$: (sparse) inferred pixel-anchor assignment

For each minibatch,

- iterate until converge: (*expit, logit*, and *exp* apply element-wise)
- E-step:

$$\Psi \propto \operatorname{expit}\left(\operatorname{logit} W + \Phi\widetilde{\Gamma^{T}} + \left(\left(X\widetilde{\Lambda^{T}}\right) \odot \Phi\right) \mathbf{1}_K \mathbf{1}_n^T\right)$$

$$\Phi \propto \exp\left(\Psi\tilde{\Gamma} + \left(X\widetilde{\Lambda^{T}}\right) \odot (\Psi 1_n 1_K^T)\right) \tag{6}$$

$$\Gamma \propto 1\alpha^T + \Psi^T$$

- M-step: Update global parameters.

$$\widetilde{\Lambda} \propto 1\eta^T + \Phi \odot (\Psi 1_n 1_K^T) \tag{7}$$

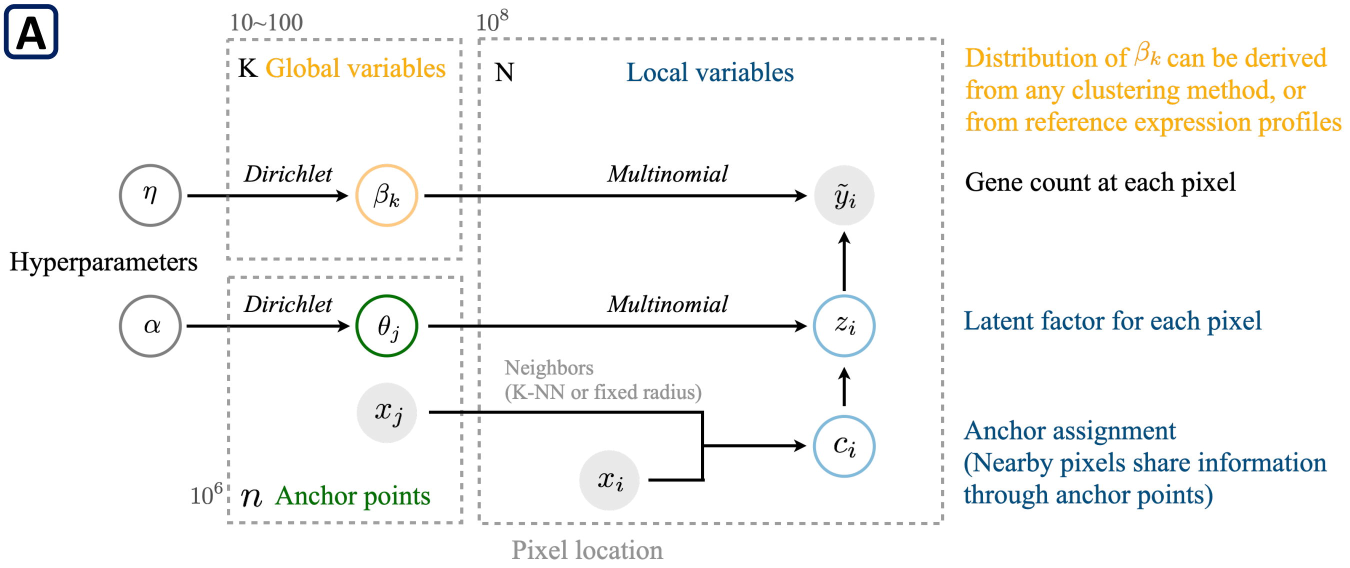
A
10~100
10$^8$
K Global variables
N
Local variables
Distribution of $\beta_k$ can be derived from any clustering method, or from reference expression profiles
Dirichlet
Multinomial
$\eta$
$\beta_k$
$\tilde{y}_i$
Gene count at each pixel
Hyperparameters
Dirichlet
Multinomial
$\alpha$
$\theta_j$
$z_i$
Latent factor for each pixel
Neighbors
(K-NN or fixed radius)
$x_j$
$c_i$
Anchor assignment
(Nearby pixels share information through anchor points)
$x_i$
10$^6$
$n$ Anchor points
Pixel location

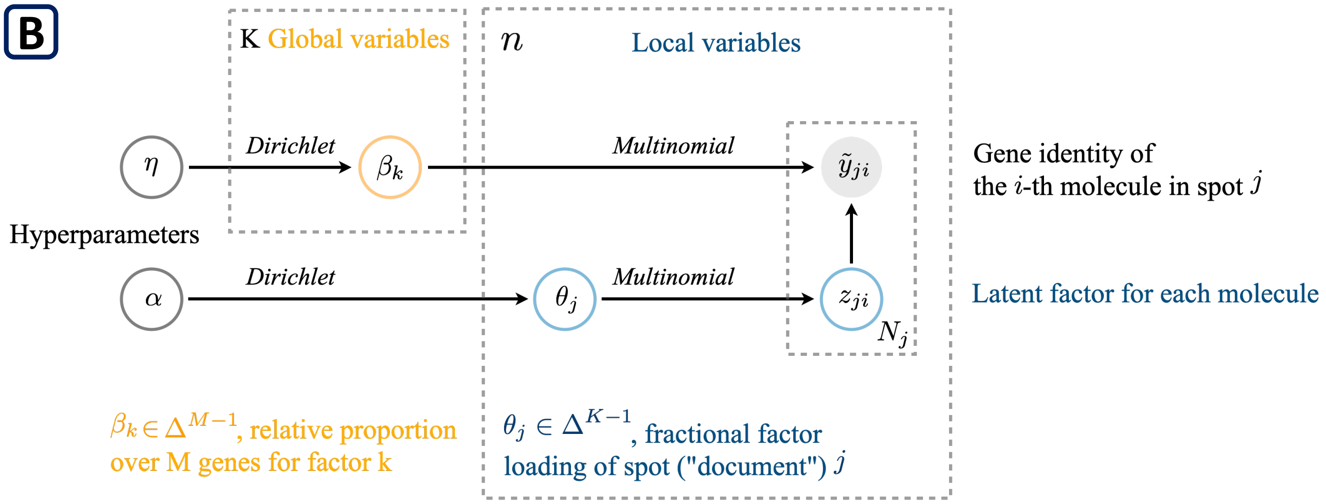
B
K Global variables
$n$
Local variables
Dirichlet
Multinomial
$\eta$
$\beta_k$
$\tilde{y}_{ji}$
Gene identity of the $i$-th molecule in spot $j$
Hyperparameters
Dirichlet
Multinomial
$\alpha$
$\theta_j$
$z_{ji}$
$N_j$
Latent factor for each molecule
$\beta_k \in \Delta^{M-1}$, relative proportion over M genes for factor k
$\theta_j \in \Delta^{K-1}$, fractional factor loading of spot ("document") $j$

**Overview of the Analyzed Datasets**

| Type | Species/ Tissue | Platform | Tissue Area (mm²) | Total Transcripts (N) | Genes Assayed (N) | Segmented Cells (N) | Methods Applied |
|---|---|---|---|---|---|---|---|
| Real | **Mouse Colon** | **Seq-Scope** | 18 | 6.8M | 27,347 | N/A | FICTURE |
| Simulated | **Mouse** | **N/A** | 2 | 8M | 500 | 8.72K | FICTURE, Baysor, GraphST |
| Simulated | **Mouse** | **N/A** | 7 | 28M | 500 | 30.49K | FICTURE, GraphST |
| Real | **Mouse E16 Embryo** | **Stereo-seq** | 115 | 700M | 30,050 | 281K | FICTURE |
| Real | **Human Breast Cancer** | **10X Xenium** | 40.2 | 42.6M | 313 | 166K | FICTURE |
| Real | **Mouse Liver** | **Vizgen MERSCOPE** | 300 | 419M | 347 | 395K | FICTURE |
| Real (Subset) | **Mouse Liver** | **Vizgen MERSCOPE** | 0.46 | 2.46M | 347 | 2.5K | FICTURE, Baysor, GraphST |
| Real | **Human Healthy Lung** | **10X Xenium** | 101 | 24M | 453 | 296K | FICTURE |

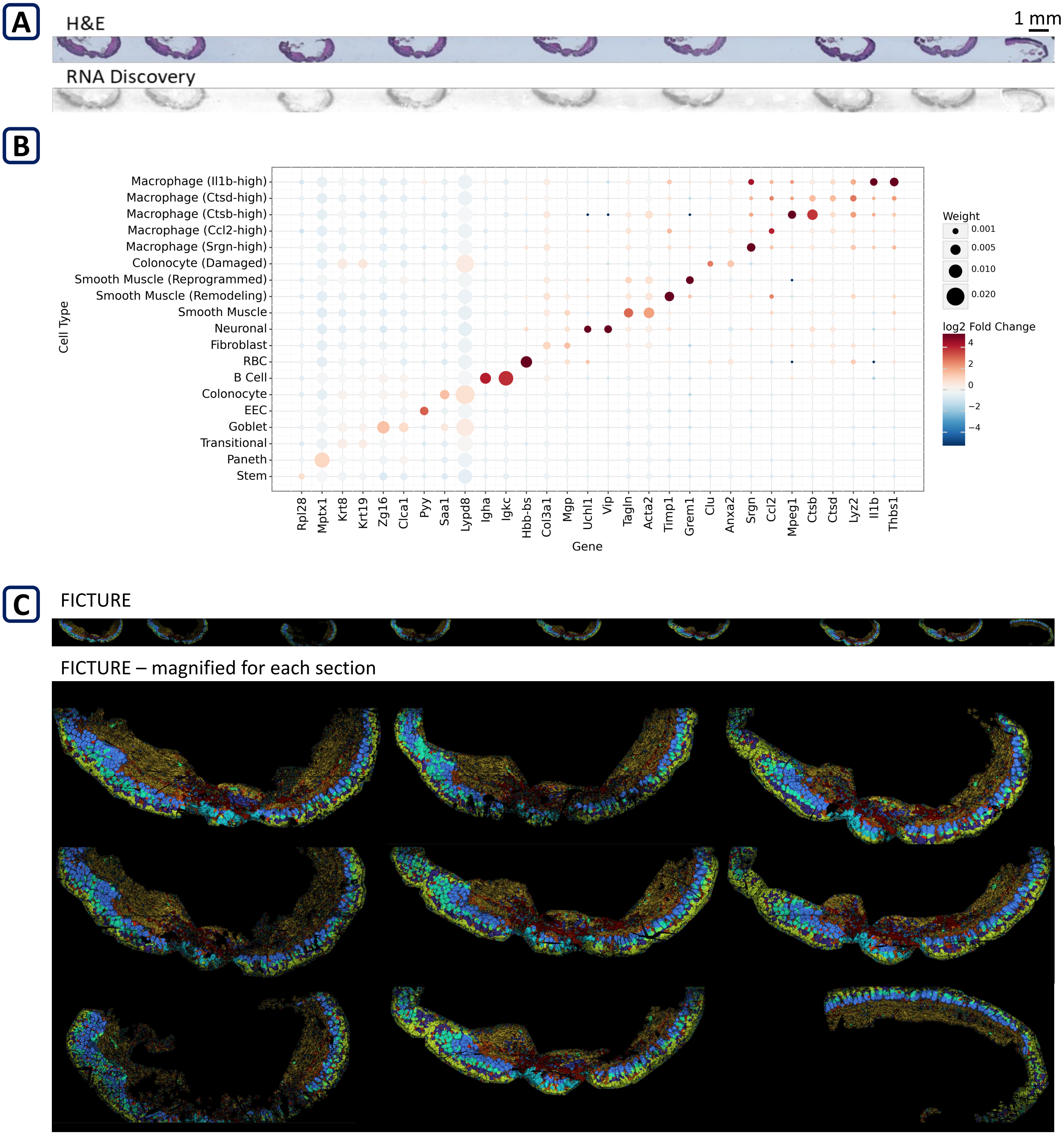
A
H&E
1 mm
RNA Discovery
B
Macrophage (Il1b-high)
Macrophage (Ctsd-high)
Macrophage (Ctsb-high)
Macrophage (Ccl2-high)
Macrophage (Srgn-high)
Colonocyte (Damaged)
Smooth Muscle (Reprogrammed)
Smooth Muscle (Remodeling)
Smooth Muscle
Neuronal
Fibroblast
RBC
B Cell
Colonocyte
EEC
Goblet
Transitional
Paneth
Stem
Cell Type
Rpl28
Mptx1
Krt8
Krt19
Zg16
Clca1
Pyy
Saa1
Lypd8
Igha
Igkc
Hbb-bs
Col3a1
Mgp
Uchl1
Vip
Tagln
Acta2
Timp1
Grem1
Clu
Anxa2
Srgn
Ccl2
Mpeg1
Ctsb
Ctsd
Lyz2
Il1b
Thbs1
Gene
Weight
0.001
0.005
0.010
0.020
log2 Fold Change
4
2
0
−2
−4
C
FICTURE
FICTURE – magnified for each section

**Supplementary Fig 4. Si et al.**

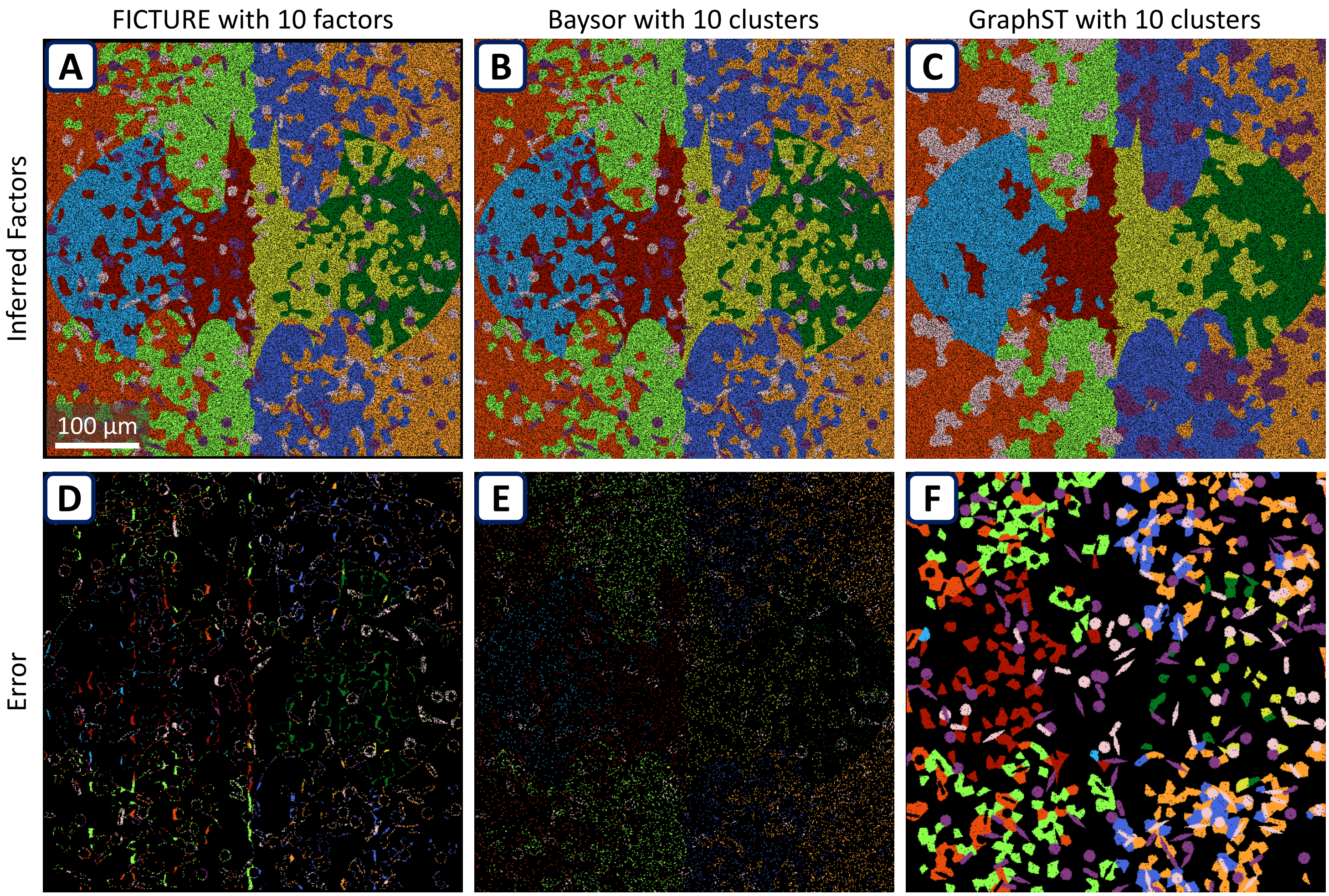

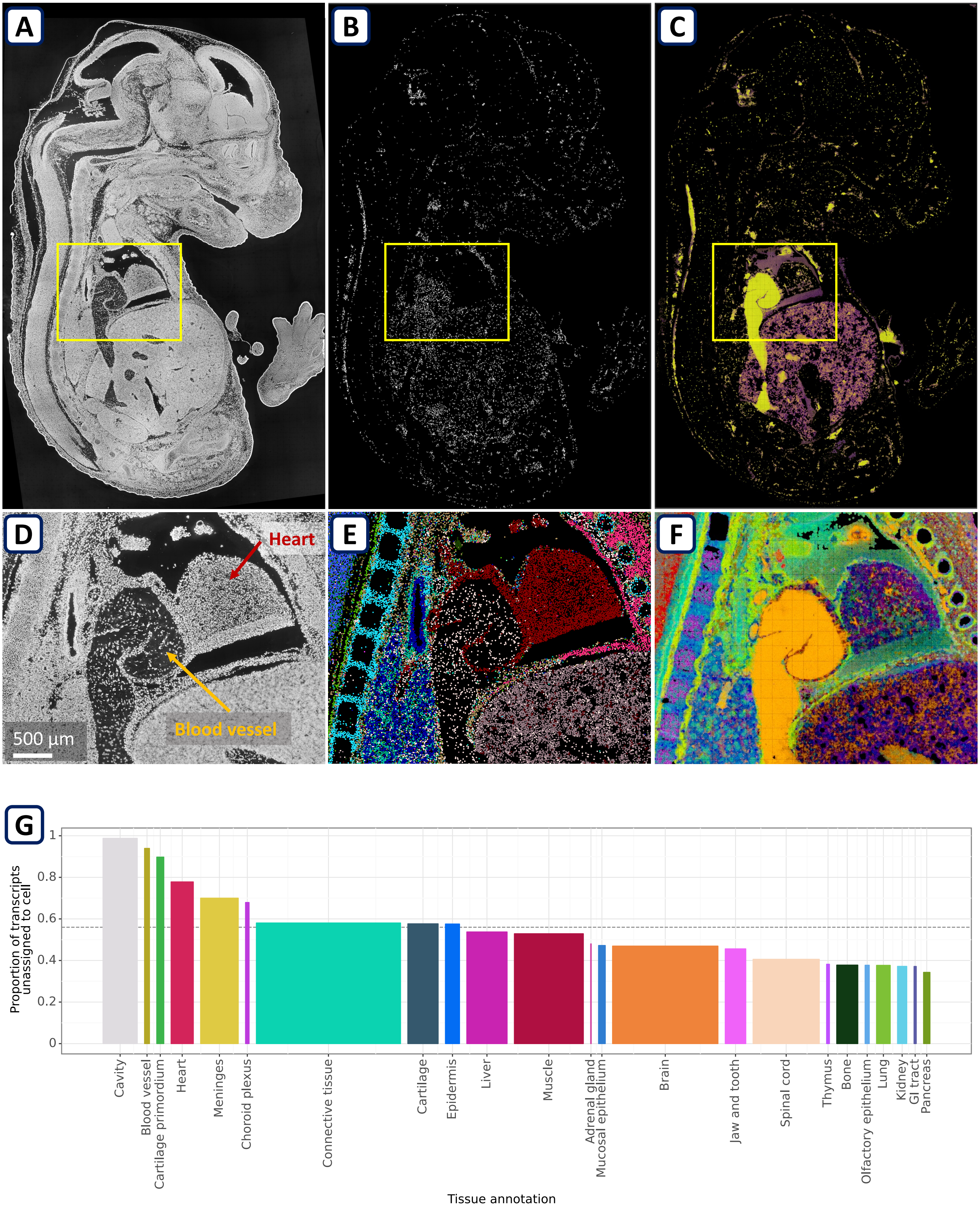
A
B
C
D
E
F
Heart
Blood vessel
500 µm
G
Proportion of transcripts unassigned to cell
1
0.8
0.6
0.4
0.2
0
Cavity
Blood vessel
Cartilage primordium
Heart
Meninges
Choroid plexus
Connective tissue
Cartilage
Epidermis
Liver
Muscle
Adrenal gland
Mucosal epithelium
Brain
Jaw and tooth
Spinal cord
Thymus
Bone
Olfactory epithelium
Lung
Kidney
GI tract
Pancreas
Tissue annotation

**Supplementary Fig 6. Si et al.**

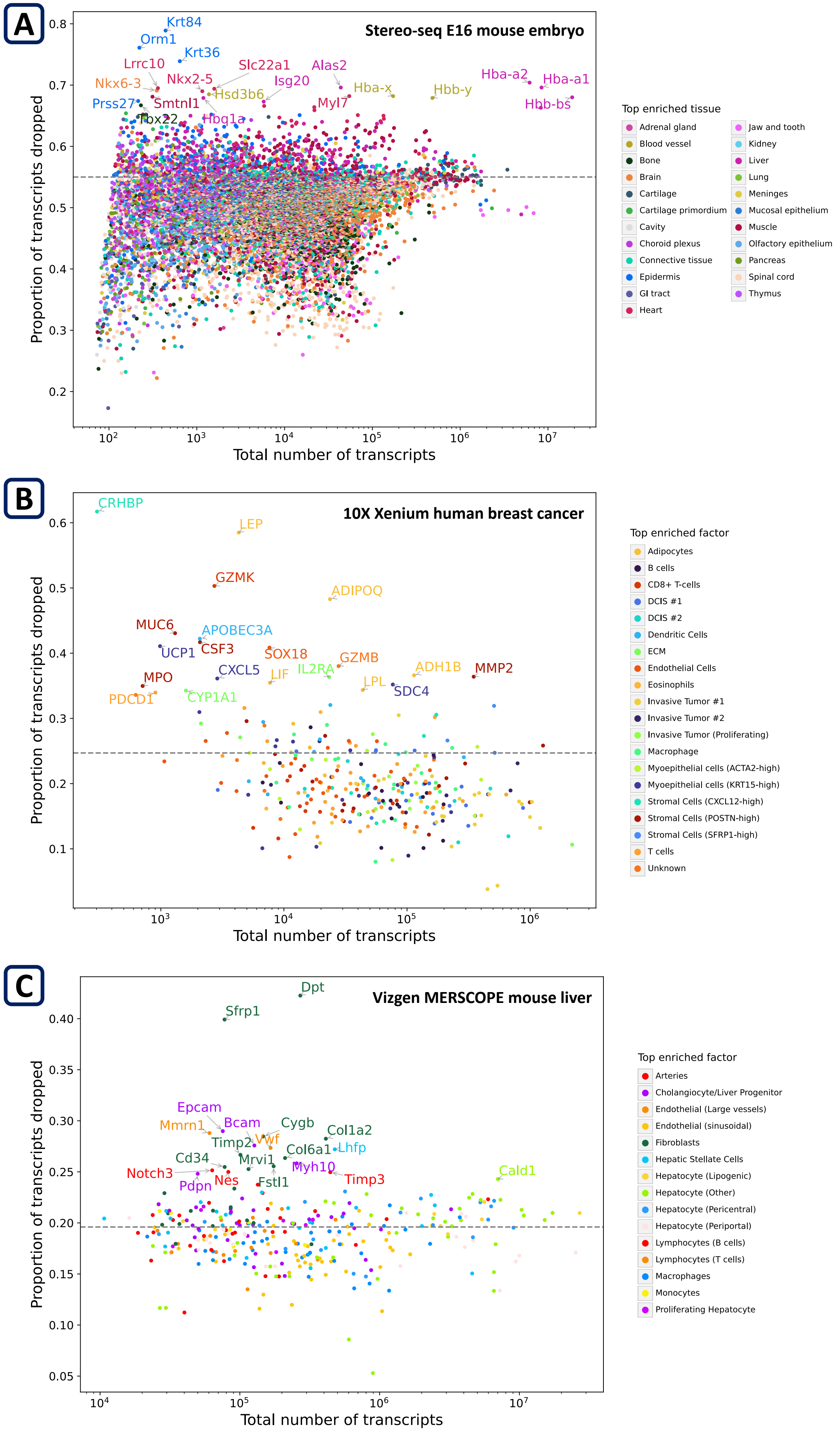

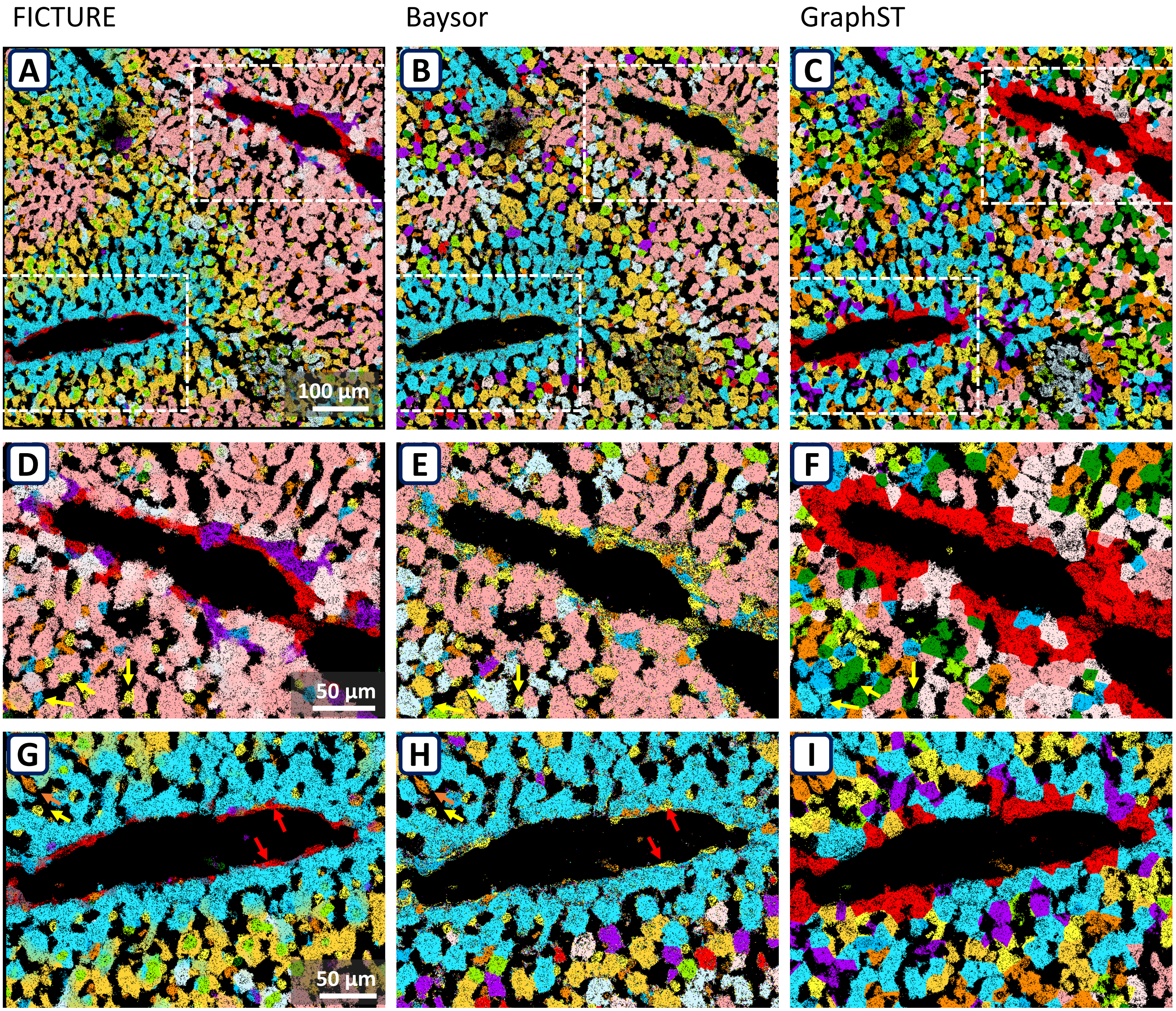
FICTURE
Baysor
GraphST
A
B
C
100 µm
D
E
F
50 µm
G
H
I
50 µm

Supplementary Fig 8. Si et al.

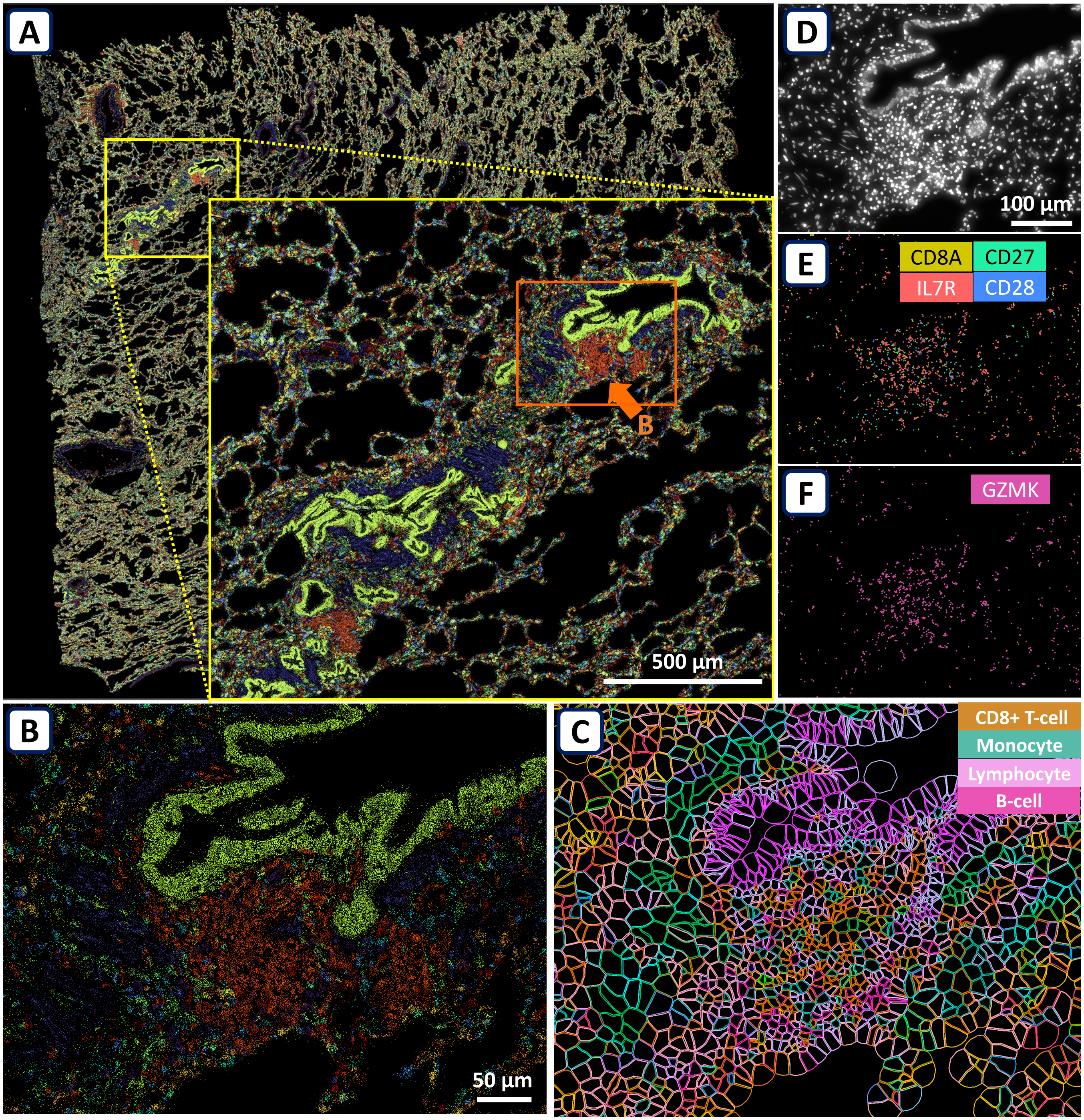
